# Single-cell RNA-seq analysis reveals lung epithelial cell-specific contributions of Tet1 to allergic inflammation

**DOI:** 10.1101/2021.12.22.473869

**Authors:** Tao Zhu, Anthony P. Brown, Lucy Cai, Gerald Quon, Hong Ji

## Abstract

Tet1 protects against house dust mite (HDM)-induced lung inflammation in mice and alters the lung methylome and transcriptome. In order to explore the role of Tet1 in individual lung epithelial cell types in HDM-induced inflammation, we established a model of HDM-induced lung inflammation in Tet1 knockout and littermate wildtype mice and studied EpCAM^+^ lung epithelial cells using single-cell RNA-seq analysis. We identified eight EpCAM^+^ lung epithelial cell types, among which AT2 cells were the most abundant. HDM challenge increased the percentage of alveolar progenitor cells (AP), broncho alveolar stem cells (BAS), and goblet cells, and decreased the percentage of AT2 and ciliated cells. Bulk and cell-type-specific analysis identified genes subject to Tet1 regulation and linked to augmented lung inflammation, including alarms, detoxification enzymes, oxidative stress response genes, and genes in tissue repair. The transcriptomic regulation was accompanied by alterations in TF activities. Trajectory analysis supports that HDM may enhance the differentiation of AP and BAS cells into AT2 cells, independent of Tet1. Collectively, our data showed that lung epithelial cells had common and unique transcriptomic signatures of allergic lung inflammation. Tet1 deletion altered transcriptomic networks in various lung epithelial cells, with an overall effect of promoting allergen-induced lung inflammation.

## INTRODUCTION

Asthma is one of the most common pulmonary disorders with high heterogeneity ^1^. Using RNA-sequencing, previous studies have noted significant transcriptomic differences in both asthmatic patients and animals with allergic lung inflammation ^2–4^. Recent studies, including ours ^2, 5–7^, have also linked DNA methylation-associated transcriptomic changes to the pathogenesis and progression of asthma. Other studies have linked enzymes involved in DNA methylation maintenance to asthma pathogenesis ^8^. Our studies in nasal epithelial samples from children and bronchial epithelial cell lines (HBECs) ^9–11^ suggest that Tet1 (Tet Methylcytosine Dioxygenase 1) may regulate childhood asthma and response to environmental exposures. Tet1 belongs to the family of ten-eleven translocation methylcytosine dioxygenases. These demethylases catalyze the hydroxylation of DNA methylcytosine (5mC) into 5-hydroxymethylcytosine (5hmC), 5-formylcytosine (5fC), and 5-carboxycytosine (5caC), interact with histone modifiers and transcription factors, and may contribute to many biological processes and responses to environmental exposures (reviewed in ^12^). In a follow up study using genetic knockout mice ^9^, we found that Tet1 plays a protective role in allergic lung inflammation. Concurrent with increased lung inflammation following house dust mite (HDM) challenges, Tet1 deficient mice showed dysregulated levels of expression at genes in several signaling pathways, including NRF2-mediated Oxidative Stress Response, Aryl Hydrocarbon Receptor (AhR) Signaling, and Interferon (IFN) signaling. These genes were also subject to Tet1 regulation in HBECs ^9^.

Given the use of RNA from total lung tissues in these previous studies, critical questions remained on the cellular origin(s) of specific inflammatory mediators and molecules following allergen challenges, and the role that Tet1 plays in regulation of these mediators/molecules in individual cell types *in vivo*. In this study, our primary goal was to explore the role of Tet1 in HDM-induced lung inflammation in individual lung epithelial cells (EpCAM^+^) from mice using scRNA-seq. The effects of Tet1 loss and HDM challenges were compared in all EpCAM^+^ cells together and in each individual cell type.

## METHODS

### Establishment of HDM-induced airway inflammation mouse model and Isolation of CD326^+^ CD31^-^ CD45^-^ (EpCAM^+^) lung epithelial cells

HDM-induced allergic inflammation was established in both Tet1^−/−^ mice (knockout/KO mice) and their littermate Tet1^+/+^ mice (wildtype/WT mice) as described in a previous study ^9^ (details in Supplementary methods and Figure S1A). All procedures on animals were approved by the Animal Experimental Ethics Committee of Cincinnati Children’s Hospital Medical Center and University of California, Davis. Airway inflammation was assessed by histology. Lobes, about 5mm large, were sectioned in half and collected from each animal. Lobes from animals of the same group were pooled into one sample. Tissues were minced in a collagenase (480U/ml)-dispase (5U/ml)-DNase (80U/ml) digestion solution in DPBS with Ca^+^ and Mg^2+^ and razor blades. The finely minced tissues were then incubated in 6mL digestion solution at 37°C for 40min with gentle inversion for 30s at 5min intervals. After incubation, samples were washed with a DPBS/1% BSA wash buffer and passed through a 70μ filter strainer. After spinning down cells and removing the supernatant, cells were treated with AKC lysis buffer to remove red blood cells and washed once more to remove dead cells and excess RNA in the buffer. CD45^+^ cells (immune cells) and CD31^+^ (endothelial cells) were then depleted using mouse CD45 and CD31 MicroBeads (Miltenyi Biotec) and the remaining cells were counted and incubated with anti-CD326-APC (eBioscience), anti-CD31-PE-cy7 (eBioscience) and anti-CD45-PE (Biolegend). CD326^+^ CD31^-^ CD45^-^ cells were collected on a MoFlo sorter (Beckman Coulter/MoFlo XDP) with >99% purity at the Research Flow Cytometry Core at Cincinnati Children’s Hospital Medical Center (Figure S1B).

### Single cell library preparation and sequencing

For 10X Genomics single cell RNA sequencing, cells were brought to a concentration of about 1000 cells/µL using the resuspension buffer. 8000 cells were targeted for cDNA preparation and put through the 10X Genomics Chromium Next GEM Single Cell 3’ v3.1 pipeline for library preparation. After cDNA preparation and GEM Generation and clean-up, cDNA was quality checked and quantified on an Agilent Bioanalyzer DNA High Sensitivity chip at a 1:10 dilution. Successful traces showed a fragment size range of approximately 1000-2000bp. 12 indexing cycles for library was used. The quality of indexed libraries were checked on an Agilent Bioanalyzer DNA High Sensitivity chip at a 1:10 dilution. Successful libraries had fragment size distributions with peaks around 400bp. Libraries were sequenced using 4 Illumina Hiseq lanes (Paired-end 75bp).

### Single-cell RNA-sequencing data analysis

We used Cell Ranger v3.1.0 ^13^ for initial processing of the scRNA-seq data. Specifically, we used Cell Ranger for sample demultiplexing, barcode processing, read alignment to the reference genome, data aggregation, and generating expression matrices. We then imported the data into Seurat v3.1.4 ^14^ for downstream processing. Cells that had fewer than 700 genes expressed, greater than 10% of reads mapping to the mitochondria, fewer than 20000 unique molecular identifiers, or that expressed Pecam1 (CD31, an endothelial marker ^15^) were filtered out. We also used cell cycle markers to identify which part of the cell cycle each cell was in during sampling ^16^, and subsequently regressed out this variable and the percentage of mitochondrial reads in each cell while normalizing and scaling the data with Seurat. In order to cluster all cell types accurately without biases from the HDM treatment, we used scAlign ^17^, a program that uses an unsupervised deep learning algorithm to correct for treatment effects that can affect cell clustering. First, the data were separated into two separate files based on treatment, and 3000 variable features were identified in each file separately. Variable features that overlapped were used for downstream clustering. We then performed single cell alignment by performing a bidirectional mapping between cells with different treatments, creating a low-dimensional alignment space where cells grouped together by type rather than by treatment ^17^. Our best results (i.e. greatest separation of clusters by type rather than by treatment) were achieved by using the first 50 principal component scores rather than individual gene level data as input to the model. The output from the scAlign model was subsequently used as input for a uniform manifold approximation and projection (UMAP) in Seurat ^14^ to visualize the cell clusters.

We then used the FindClusters function from Seurat (a shared-nearest-neighbor-based approach) to cluster the cells, using a wide range of resolutions from 0.01 to 2 to identify the best resolution for our dataset. FindClusters is a shared nearest neighbor modularity optimization based clustering algorithm. Our eventual selection was a resolution of 1, yielding 11 initial cell clusters. We then used markers from the single-cell mouse cell atlas ^18^ and another study on lung epithelial cells ^19^ to identify cell clusters. For each cluster, we plotted the gene expression levels of the markers from each epithelial cell type in our references, and identified which cell type each cluster represented. We merged some of the original 11 clusters identified by Seurat ^14^ that expressed similar markers, only merging sister nodes in the cell cluster tree generated by BuildClusterTree from Seurat. After merging similar clusters, we had a total of nine different labeled cell clusters in our dataset. We then used Seurat to calculate cluster means and perform differential expression analyses via the Wilcox Rank Sum Test. For each cell type individually as well as for all cells combined, we performed five differential expression comparisons: 1) WT-HDM vs. WT-Saline, 2) KO-Saline vs. WT-Saline, 3) KO-HDM vs KO-Saline, 4) KO-HDM vs. WT-HDM and 5) KO-HDM vs. WT-Saline. Genes that were differentially expressed had a false discovery rate (FDR) of 0.05 or less, and a fold change of at least 1.2. Genes also had to be expressed in at least ten percent of one of the comparison groups to qualify as differentially expressed. Pathway enrichment analyses for differentially expressed genes from each comparison were performed using IPA (QIAGEN Inc., https://www.qiagenbioinformatics.com/products/ingenuitypathway-analysis).

### Transcription factor activity analysis

We used DoRothEA ^20–22^ to assess the relative transcription factor (TF) activity levels in different cell types and/or treatments. DoRothEA computes activity levels for transcription factors using the expression levels of targets of each TF instead of the expression level of the TF itself. For our analysis, we chose to include only highly confident interactions between TFs and target genes (confidence levels “A”, “B”, or “C”). We used VIPER for statistical analysis of the TF activity levels, as VIPER considers the mode of each TF-target interaction and has been shown to be appropriate for single-cell analysis ^22, 23^. We compared TF activity between cell types and plotted top 50 by activity level variance. We also compared TF activity within cell types by sample type in AT2 and ciliated cells and plotted top 25.

### Single-cell trajectory analysis

In order to assess the differentiation of AP and BAS cells to AT2 cells, we performed single-cell trajectory analysis using Monocle ^24^. First, the dataset was subsetted to include only AP, BAS, and AT2 cells using Seurat ^14^. Next, cells were ordered based on the progress they had made through the differentiation process via reversed graph embedding using Monocle, then dimensionality reduction was performed using the “DDRTree” reduction method, and cell trajectories were visualized. We also performed trajectory analyses on different subsets of AP, BAS, and AT2 cells to better understand how HDM treatment and Tet1 deletion impacted the differentiation of AP and BAS cells into AT2 cells. We performed cell trajectory analyses on 1) all AP, BAS, and AT2 cells; 2) only wildtype AP, BAS, and AT2 cells; 3) only AP, BAS, and AT2 cells that had Tet1 knocked out; 4) only saline treated AP, BAS, and AT2 cells; 5) only HDM treated AP, BAS, and AT2 cells. For each of these comparisons, we also performed branched expression analysis modeling within Monocle to identify genes that were differentially expressed along the trajectory at branch points that roughly separated AP, BAS, and AT2 cells. However, this was not possible in the saline cell only group, as there were no branch points that clearly separated these different groups of cells. We then used Monocle to make heatmaps visualizing the expression patterns of genes that showed a significant change in expression along these trajectories (q < 1 x 10^-4^).

## RESULTS

### Single-cell analysis identified nine cell types in the EpCAM^+^ CD45^-^ CD31^-^ cells from mouse lungs

In order to understand the contribution of Tet1 expression in lung epithelium to allergic inflammation, we isolated EpCAM^+^ CD45^-^ CD31^-^ (EpCAM^+^) lung epithelial cells from animals in an established HDM-induced allergic airway inflammation model ^9^ and characterized them using single-cell RNAseq (overall study design in Figure 1A, treatment protocol and isolation strategy in Figure S1A and S1B). To demonstrate the successful establishment of allergen-induced lung inflammation, pathological alterations were observed. Consistent with our previous observations ^9^, airway inflammatory cells infiltration, goblet cell proliferation, smooth muscle hypertrophy, and airway basement membrane thickening were observed in HDM-challenged WT (WT-HDM) mice included in the single-cell analysis (Figure S1C and S1D). Slightly more inflammatory cells infiltration, goblet cell proliferation, and airway mucus secretion were observed in HDM-challenged KO mice (KO-HDM) included in this scRNA-seq analysis. No pathological changes were observed in both WT-Saline and KO-Saline mice.

**Figure 1.**
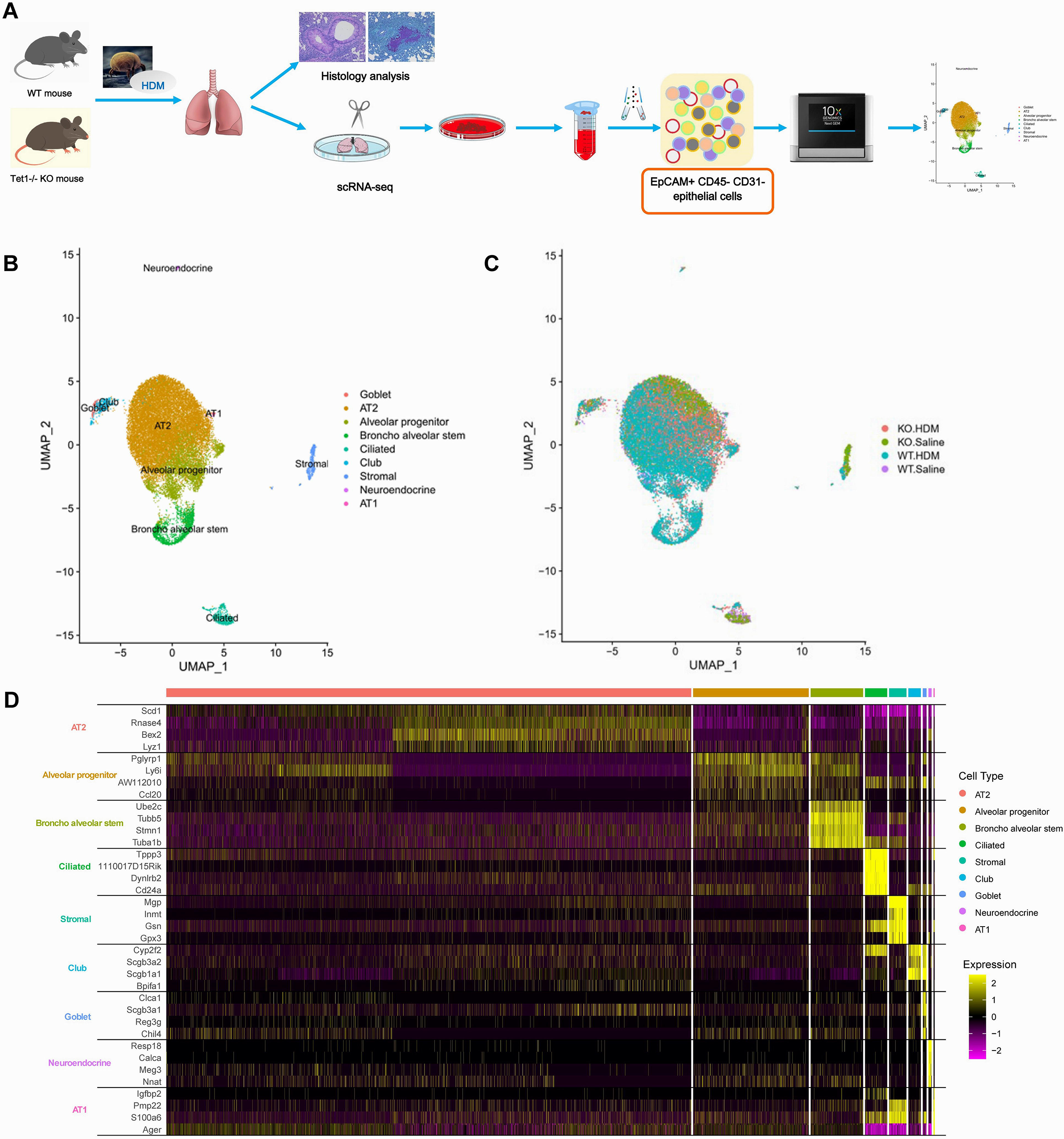
Single-cell RNA sequencing clustering analysis identifies 9 cell types in EpCAM^+^ CD45^-^ CD31^-^ lung cells. (A) Schematic overview of the study workflow. (B) UMAP visualization of clustering revealed 9 cell populations. Population identities were determined based on marker gene expression. (C) UMAP visualization of EpCAM^+^ CD45^-^ CD31^-^ lung cells in Tet1 KO mice and WT mice with and without HDM challenge. (D) Heatmap of top 4 markers in 9 cell clusters.

Among the 25,437 cells we sequenced that passed our quality filters, we identified nine distinct cell types (Figure 1B/1C) based on established marker expression patterns (Figure 1D, Table S1). In order of abundance, we identified AT2 cells, alveolar progenitor (AP), broncho alveolar stem (BAS), ciliated cells, stromal cell, club cells, goblet, neuroendocrine cells, and AT1 cells (Figure 2 and Table S2). AT2 cells were by far the most abundant (17,729 out of 23,437 were AT2 cells). On average, we observed a 24.9% increase in AP cells (2.0% ± 0.6% 26.9% ± 2.1%) and 11.1% increase in BAS cells (1.0% ± 0.2% 12.1% ± 3.0%) following HDM challenges, while the percentage of AT2 cells (85.3% ± 1.4% 56.1% ± 0.04%) and ciliated cells (4.6% ± 1.5% 1.5% ± 0.7%) decreased following HDM challenges (Figure 2 and Table S2). A small increase in the percentage of goblet cells following HDM challenges was also observed, consistent with goblet cell metaplasia in histological studies (Figure S1C and S1D). Although we expect cell loss during single-cell suspension preparation and cell sorting, these results suggest dynamic changes of AP, BAS, AT2, ciliated and goblet cells in HDM-induced lung inflammation. Additionally, since stromal cells are not epithelial cells, we will not discuss DEGs in stromal cells further in this study.

**Figure 2.**
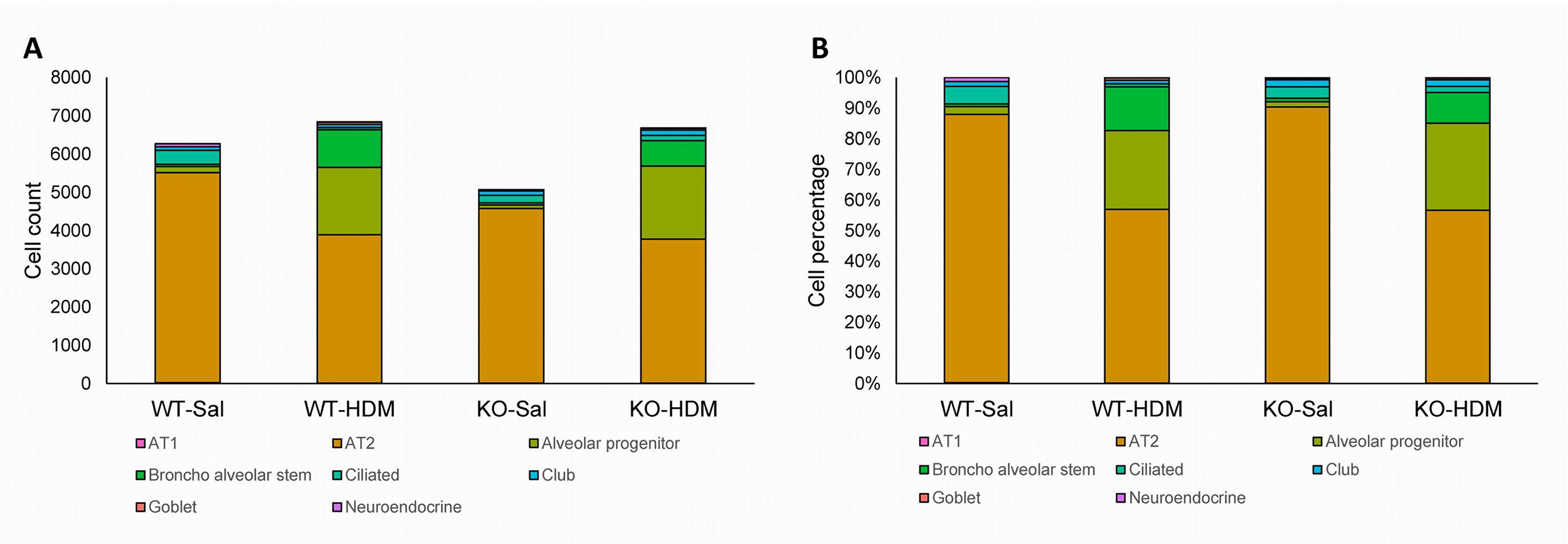
The abundance and percentage of each cell type. (A) Abundance of each cell type at each condition; (B) Percentage of each cell type at each condition

### Exposure to HDM led to cell type-specific changes in the EpCAM^+^ lung epithelial cells

We identified 830 differentially expressed genes (DEGs) between all WT-HDM and WT-Sal cells in our analysis of all EpCAM^+^ cells (hereafter referred to as bulk cell analysis), regardless of cell type (Table 1, Table S3 and Figure S2). Of the 830 DEGs, 560 were upregulated and 270 were downregulated in WT-HDM. As predicted, several pathways were significantly activated or inhibited, including Oxidative Phosphorylation (activation z-score=7.353), Sirtuin Signaling Pathway (z-score=-2.897) and Xenobiotic Metabolism AHR Signaling Pathway (z-score=-2.111) (Table S4). We further identified DEGs within each of the 8 cell types in the WT-HDM vs WT-Saline comparison. In total, we found 1773 DEGs, greater than the number of DEGs identified in the bulk cell analysis (Table 1). All cell types except AT1 cells had some differential expression associated with HDM treatment. Of these 1773 DEGs, 616 were unique to one cell type (Figure 3). AT2 cells had the most differentially expressed genes associated with HDM treatment (n=732), followed by AP (n=431) (Table 1, Figure 3A, Table S5, and Figure S2). Among the 732 DEGs identified in AT2 cells, 476 were upregulated (including *Il33*, *Gstt1*, *Isg15*, *Iftm3*, and *Cxcl15*) and 256 were downregulated (e.g., *Gstm1*) in WT-HDM compared to WT-Sal. Many genes, including *Il33* ^25^, *Cxcl5* ^26^, *Cxcl15* ^27^, *Ccl20* ^28^, *Cxcl3* ^29^, *Cxcl1* ^30^, *C3* ^31^, that have been linked to asthma pathogenesis showed cell type-specific response to HDM (Figure 4F-4H, Tables S5 and S6). Subsequently, pathway analysis identified a total of 120 significantly enriched pathways (Table S7). Besides 98 pathways shared with the bulk cell analysis, 22 pathways were unique to AT2 cells, including Ferroptosis Signaling Pathway, Chemokine Signaling, LPS-stimulated MAPK Signaling, etc. Additionally, 79 IPA pathways were enriched in the 318 unique DEGs in AT2 cells.

**Figure 3.**
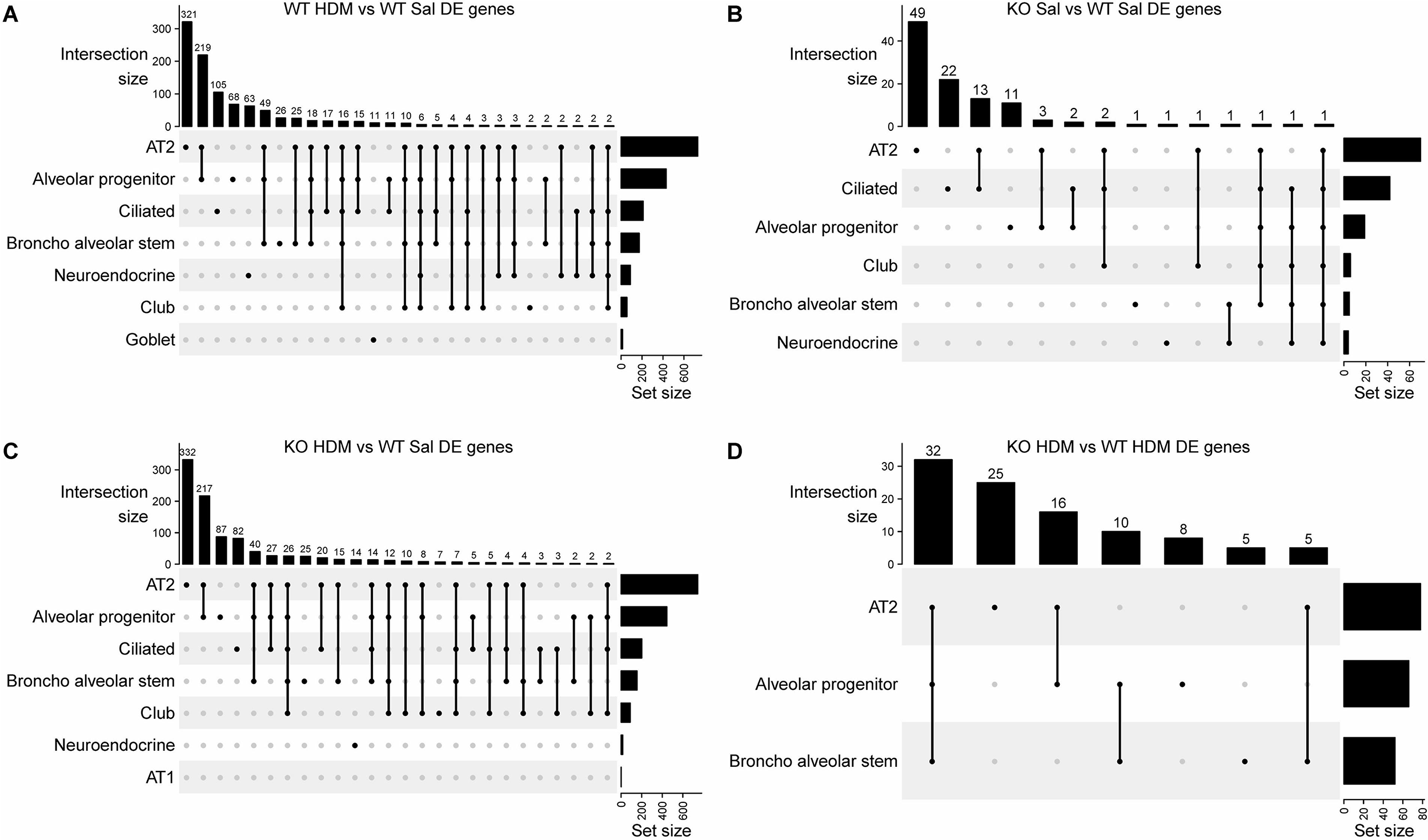
UpSet plots for overlapped DEGs among 8 individual cells types. (A) WT-Sal vs WT-HDM; (B) WT-Sal vs KO-Sal; (C) KO-HDM vs WT-Sal; (D) KO-HDM vs WT-HDM. Dots below vertical bars represent overlapped DEGs among different cell types. Horizontal bars represent the total number of DEGs in each cell types.

**Figure 4.**
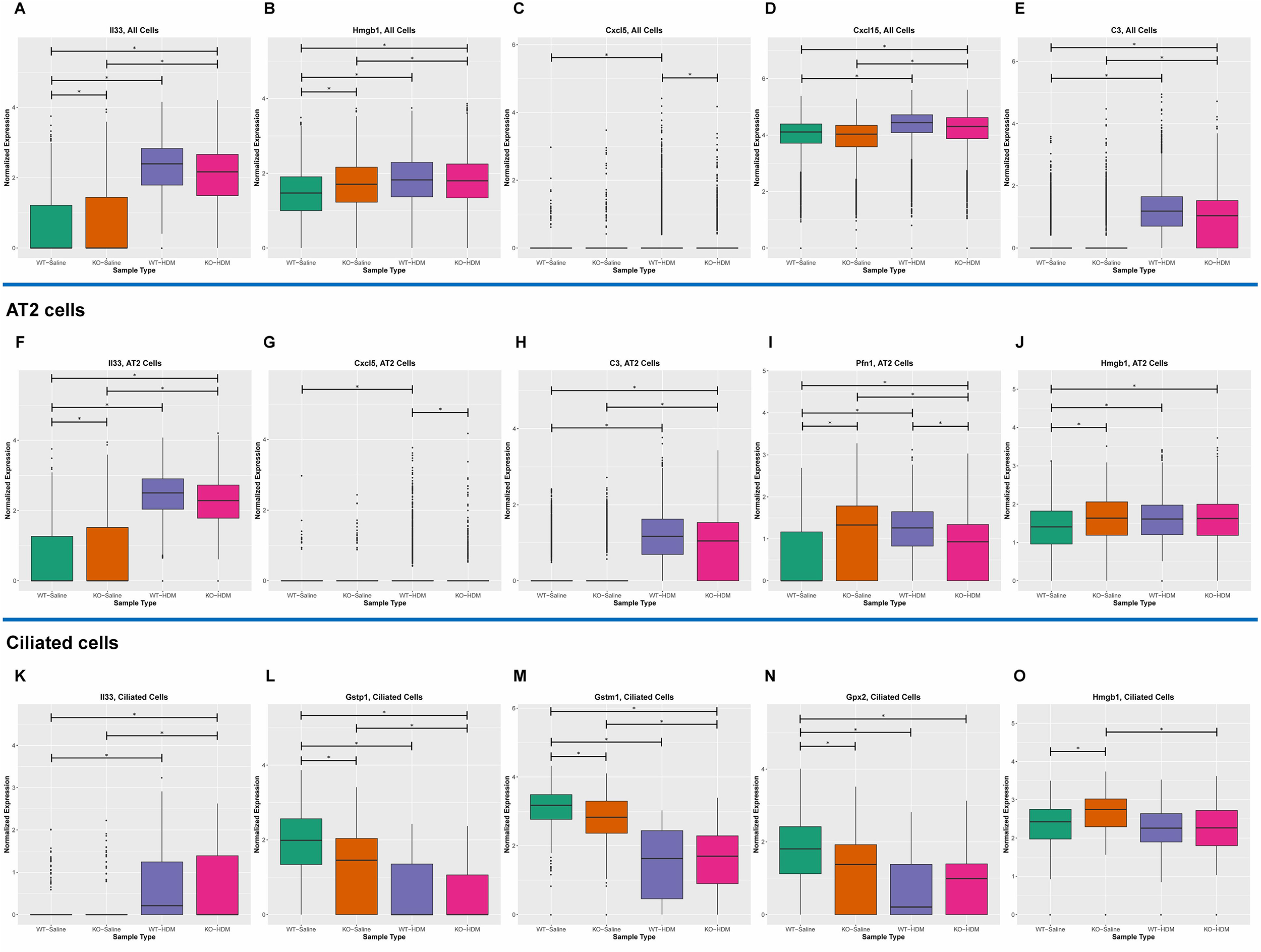
Cell-type specific changes resulting from HDM challenges and/or Tet1 deficiency. (A-E) All EpCAM^+^ cells; (F-J) AT2 cells; (K-O) Ciliated cells.

**Table 1.**
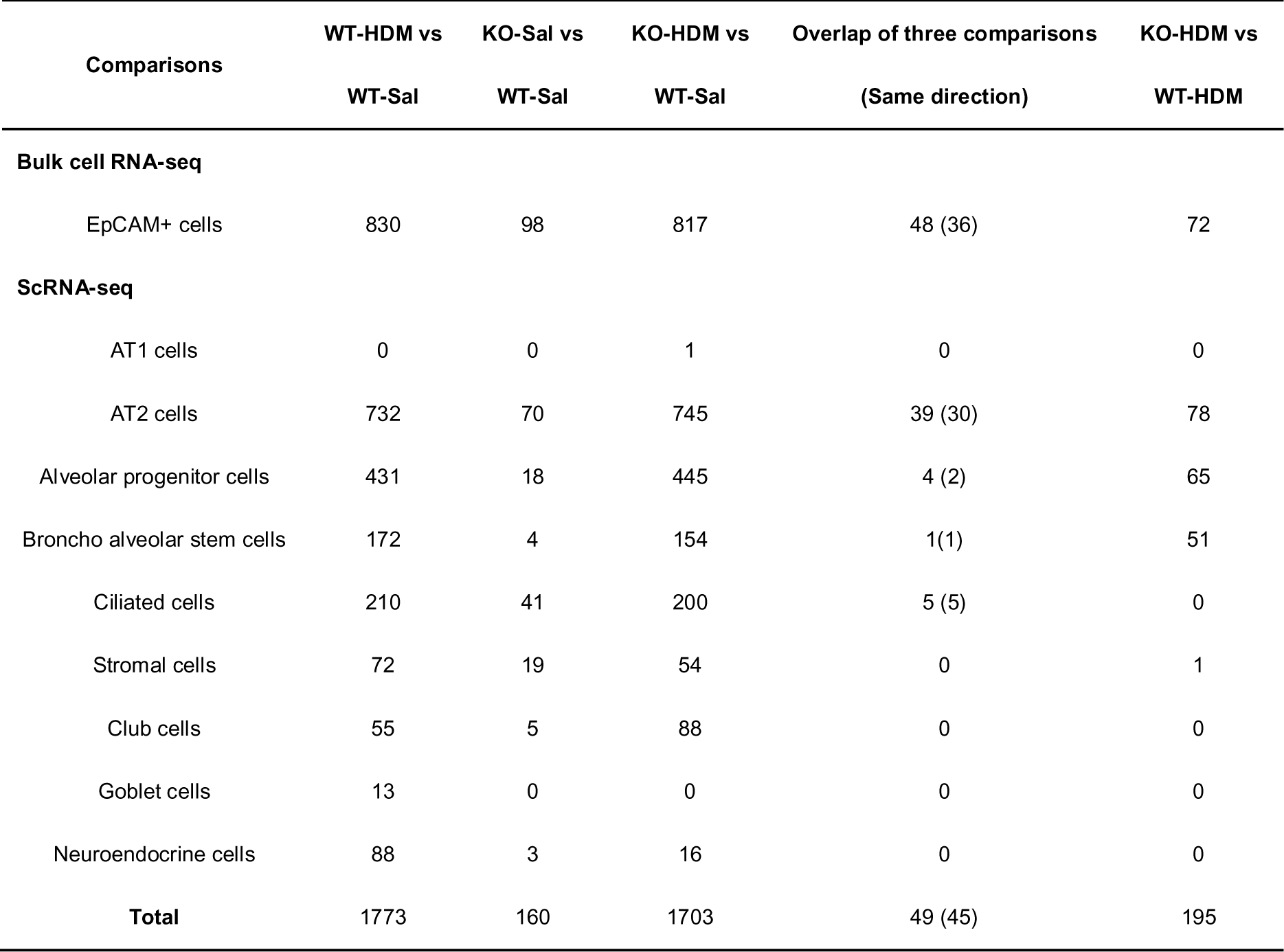
Number of Differential expression genes (DEGs) of EpCAM^+^ cells.

Chronic airway inflammation induced by inhaled allergens plays a critical role in the pathophysiology of asthma. Ciliated cells are one major cell type that contributes to mucociliary clearance of pathogens, and therefore is subject to injury and chronic inflammation. Inflammatory mediators, such as Th2 cytokines (IL-4, IL-5, and IL-13), can directly cause inflammation and injury in ciliated cells in asthma ^32, 33^. Moreover, injured ciliated cells will release more inflammatory mediators, such as TSLP and IL-33, leading to more airway inflammation and obstruction ^34, 35^. A total of 210 DEGs were found in ciliated cells in the WT-Sal vs WT-HDM comparison, with 104 of these genes being unique to ciliated cells (Table 1, Figure 3A, Figure S2, Tables S5 and S6). Compared with WT-Sal, 155 DEGs were upregulated (e.g., *Il33* and *Ifitm3*) and 55 DEGs were downregulated (e.g., *Gstp1*, *Gstm1* and *Gpx2*) in WT-HDM ciliated cells (examples shown in Figure 4K-4N). Other DEGs from this comparison, such as *Fos* ^36^, *Jun* ^37^, *Hes1* ^38^, *Cd14* ^39^ and *C3* ^31^, have been associated with asthma pathogenesis. Subsequently, 96 significantly enriched pathways in these DEGs were identified (Table S8). NRF2-mediated Oxidative Stress Response was predicted to be inhibited (z-score=-2.333). Furthermore, 18 IPA pathways were enriched in the 104 unique DEGs in ciliated cells (Table S8). Among them, some pathways, such as NRF2-mediated Oxidative Stress Response (*Dnajc5*, *Gpx2*, *Gstm6*, and *Maff*) and Antigen Presentation Pathway (*Cd74* and *Hla-e*) ^40^, have been linked to asthma. Collectively, these results suggest that different types of EpCAM^+^ lung epithelial cells contribute to the allergic responses to HDM and pathogenesis of asthma, with common features as well as unique functions.

### Tet1 knockout led to cell-specific changes in baseline EpCAM^+^ lung epithelial cells

In the bulk analysis comparing KO-Sal and WT-Sal cells, 98 genes were differentially expressed (Table 1, Figure S2, and Table S3). There were 27 significantly enriched pathways associated with these DEGs (Table S4), including several stress-related pathways (Protein Ubiquitination Pathway/Unfolded protein response/eNOS Signaling/NRF2-mediated Oxidative Stress Response/Glutathione Redox Reactions I), and pro-inflammatory pathways (Interferon Signaling/IL-6 Signaling/ IL-12 Signaling and Production in Macrophages). Furthermore, we identified DEGs in each cell type (Table 1 and Table S5). A total of 70 DEGs were found in AT2 cells, including 48 DEGs unique to AT2 cells (e.g., *Il33*, *Malat1* and *S100a6*) (Table S5 and Figures 3B, example plots shown in Figures 4, 5 and Figure S3). Compared with WT-Sal, 42 DEGs were upregulated (e.g., *IL33*) and 28 DEGs were downregulated (e.g., *Lrg1* and *Areg*) in KO-Sal. There were 45 significantly enriched IPA pathways (Table S7), including 17 found in bulk cell analysis and 28 unique ones to AT2 cells (e.g., AhR Signaling/Xenobiotic Metabolism AHR Signaling Pathway). Several genes, *Areg* ^41^ and *Bpifa1* ^42^, *Hspa8* ^43^ and *Lgals3* ^44^ (also DEGs in ciliated cells), have previously been linked to asthma.

**Figure 5.**
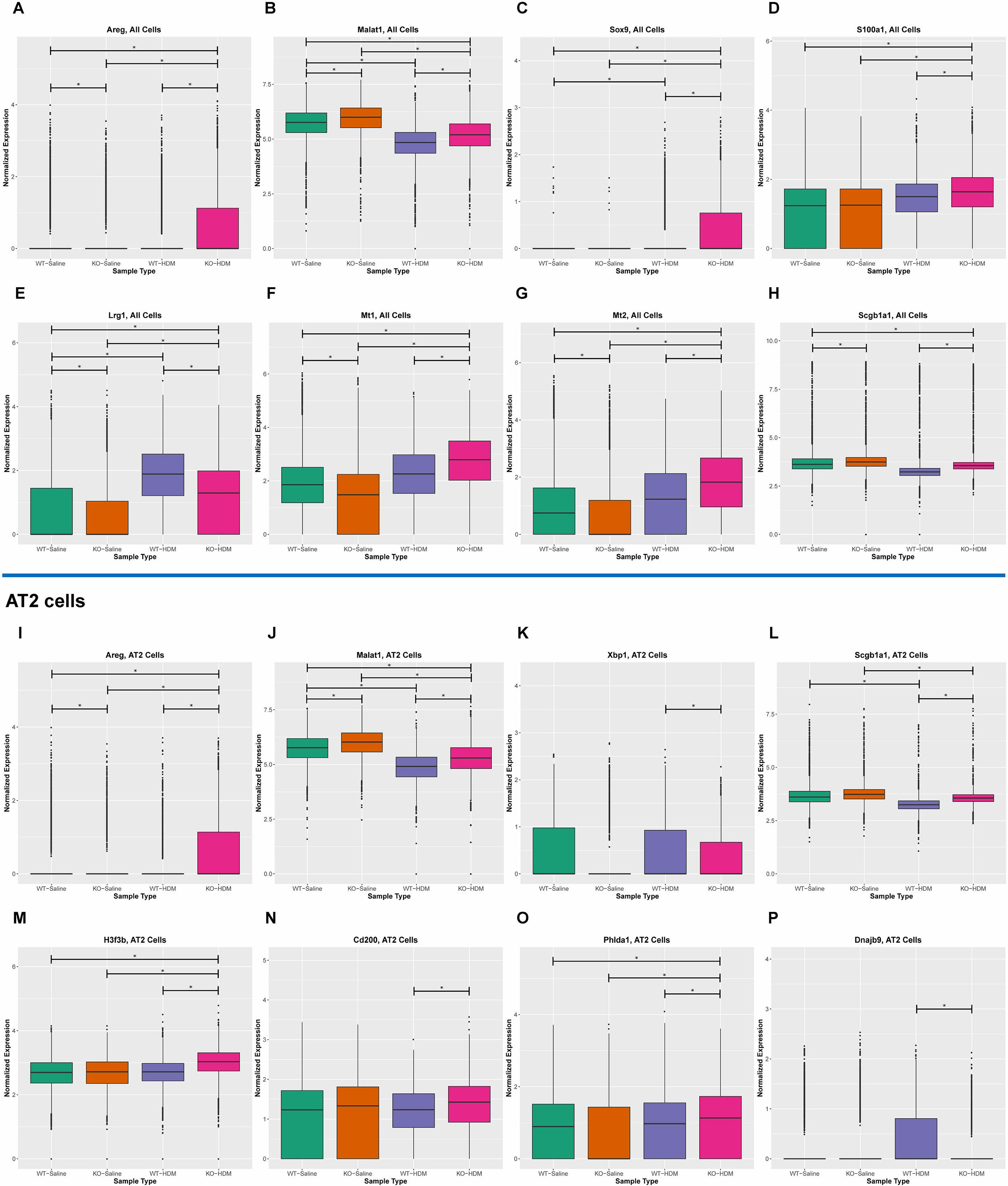
Tet1 loss promotes HDM-induced lung inflammation by upregulation of proinflammatory responses and tissue damage and repair, and downregulation of genes involved in oxidative stress. (A-H) All EpCAM^+^ cells; (I-P) AT2 cells.

Additionally, 41 Tet1 loss-induced DEGs were found in ciliated cells (Table 1, Table S5 and Figure 3B). Compared with WT-Sal, 24 DEGs were upregulated and 17 DEGs were downregulated in KO-Sal. Several asthma-related genes were identified: detoxification enzymes (*Cyp2a5* ^45^, *Gsto1* ^46^, *Gstp1* ^47^, *Gstm1* ^48^, and *Aldh1a1* ^46^) were downregulated while *Hmgb1* (alarmin) ^49 50^ was upregulated (Figure 4L-4O Tables S5 and S6). Consistent with our observations following HDM treatment, the downregulation of the detoxification enzymes following Tet1 KO (*Gstp1, Gstm1*, and *Gpx2*) were only found in ciliated cells (Figure 4L-4N and Table S6). *Hmgb1* was also upregulated in AT2 cells following Tet1 KO. 49 significantly enriched pathways were identified in ciliated cells (Table S8), which revealed the significant inactivation of the Xenobiotic Metabolism AHR Signaling Pathway (z-score= -2.449) and enrichment of several other stress response pathways.

In summary, our single-cell analysis comparing WT-Saline and KO-Saline found that loss of Tet1 in saline-treated mice led to 1) increased expression of alarmins, particularly *Il33* (AT2 cells) and *Hmgb1* (AT2 and ciliated cells), that promote type 2 inflammation, 2) upregulation of genes that promote proliferation and migration of airway smooth muscle cells and tissue remodeling [*Malat1* (AT2 cells) and *Pfn1* (6 cell types)], and 3) reduced expression of detoxification enzymes (*Gpx2*, *Gstm1*, *Gsto1*, and *Gstp1*) in AhR Signaling in ciliated cells. Together, this gene dysregulation following Tet1 loss may have resulted in a pro-inflammatory state in AT2 and ciliated cells.

### Tet1 knockout and HDM exposure induced overlapping cell type-specific changes in EpCAM+ lung epithelial cells

To further explore the effects of Tet1 knockout in HDM-induced lung inflammation, we searched for overlapping genes among the WT-HDM vs WT-Sal, KO-Sal vs WT-Sal, and KO-HDM vs WT-Sal comparisons. 48 overlapping DEGs were identified, including 36 with changes in the same directions (14 upregulated and 22 downregulated) in bulk cell analyses (Table 1 and Table S9). Thirteen genes were linked to different aspects of asthma, including 8 upregulated genes (*Il33, Hmgb1*, *Ager*, *Chia1*, *Ifitm3, Igfbp7*, *Pfn1*, *Retnla*) and 5 downregulated genes (*Atf3*, *Btg2*, *Klf6*, *Neat1*, and *Sec14l3*) (Figures 4A/4B and S3-B1-B12). Eight mitochondrially encoded genes were also among the 36 overlapping DEGs, but their role in asthma is not clear. Next, overlapping DEGs within each cell type were identified and investigated further. In AT2 cells, 39 genes were found differentially expressed in all 3 different comparisons (WT-HDM vs WT-Sal, KO-Sal vs WT-Sal, and KO-HDM vs WT-Sal) in AT2 cells (Table S9). 30 genes had changes in the same direction (12 upregulated and 18 downregulated), 27 of which were also overlapping in the bulk cell analyses. Meanwhile, there were 5 genes that were consistently downregulated in ciliated cells among the three aforementioned comparisons (Table S9), including asthma-associated genes, *Gstp1*, *Gstm1*, and *Gpx2* (Figure 4L-4N). Additionally, we found that 2 genes (*Dmkn* and *Chia1*) in AP cells and 1 gene (*Nnat*) in BAS cells were consistently changed in these 3 comparisons. IPA analysis revealed several significant diseases and functions affected by these overlapping genes, including inflammation of organ in EpCAM^+^ cells and AT2 cells (Figure 6A/6B), cell proliferation of fibroblast in AT2 cells (Figure 6C), and synthesis of reactive oxygen species in ciliated cells (Figure 6D).

**Figure 6.**
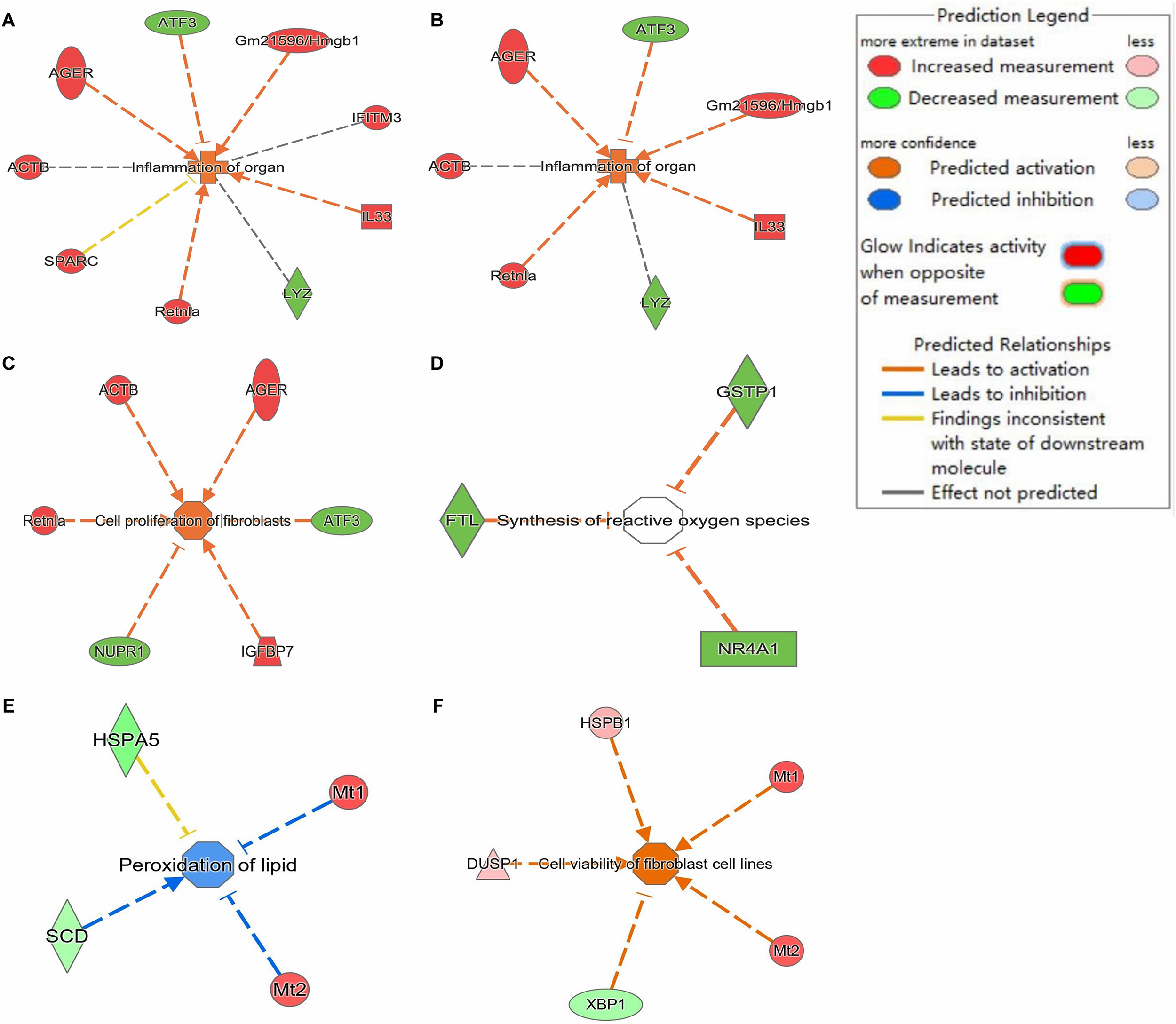
Diseases and Function analysis in EpCAM^+^, AT2 and ciliated cells via IPA. (A-D) DEGs with consistent direction changes among 3 comparisons (WT-HDM vs WT-Sal, KO-Sal vs WT-Sal, and KO-HDM vs WT-Sal). (A) Inflammation of organ in EpCAM^+^ cells. (B) Inflammation of organ in AT2 cells. (C) Cell viability of fibroblast in AT2 cells. (D) Synthesis of reactive oxygen species in ciliated cells. (E) Peroxidation of lipid and (F) Cell viability of fibroblast cell lines in DEGs in KO-HDM vs WT-HDM in AT2 cells.

The finding that several genes had consistent gene expression changes following Tet1 knockout, HDM exposure, and Tet1-knockout plus HDM suggests that this cluster of genes contributed substantially to Tet1 knockout-enhanced HDM-induced lung inflammation in mice. Furthermore, our data suggest that the effects of Tet1 knockout in allergen-induced lung inflammation were cell-type dependent, and that individual cell types play specific and sometimes separate roles in the pathogenesis of asthma.

### Tet1 dysregulates the single-cell transcriptomic signature of HDM-induced lung inflammation in mice

In order to further understand the effects of Tet1 loss in presence of HDM, we looked into the 72 genes that were differentially expressed in KO-HDM compared with WT-HDM in all EpCAM^+^ cells (Figure S2 and Table S3). 34 genes were upregulated [e.g., *Areg* (Fig 5A), *Malat1* (Fig 5B), *Sox9* (Fig 5C), and *S100a1* (Fig 5D)] and 38 were downregulated [e.g., *Lrg1* (Fig 5E) *and Ifitm3* (Fig S3-B3), *Cxcl1*, and *Rnase4*] (Table S6). Among the 72 DEGs, 29 significantly enriched pathways were identified (Table S4), including several stress response pathways (e.g., Protein Ubiquitination Pathway/ Unfolded protein response/ Endoplasmic

Reticulum Stress Pathway/ NRF2-mediated Oxidative Stress Response) and proinflammatory pathways (e.g., Interferon signaling pathway/ Antigen Presentation Pathway/ Natural Killer Cell Signaling). This is consistent with our previous observations in bulk RNA-seq from total lungs in mice challenged using the same exposure protocol ^9^. Surprisingly, several genes upregulated by HDM challenges, such as *Cxcl5* (Fig 4C)/*Isg15*/*Ifitm3* (Fig S3-B3)/*Lrg1* (Fig 5E), showed reduced expression in KO-HDM compared to WT-HDM (Table S6). In contrast, other genes involved in tissue injury/repair and remodeling, such as *Areg* (Fig 5A)^51, 52^, *Malat1* (Fig 5B) ^53, 54^, *S100a1* (Fig 5D) ^55, 56^, *Mt1/2* (Fig 5F/5G) ^57, 58^, *Scgb1a1* (Fig 5H) ^59, 60^ and *Clu* ^61, 62^, were upregulated in KO-HDM compared to WT-HDM, even though some of these genes (e.g., *Areg*) were downregulated in KO-Sal mice compared to WT-Sal mice.

We then compared KO-HDM with WT-HDM in each cell type. Interestingly, of the 195 DEGs identified across all cell types, 194 were identified in AP cells, AT2 cells, or BAS cells (Table 1 and Figure 3D), and 32 DEGs were shared among all three of these cell types. All 32 of these common DEGs changed in the same direction: 16 were upregulated (e.g., *Areg*, *Malat1*, *Mt1*/*2*, and *Scgb1a1*) and 16 were downregulated (e.g., *Cxcl5*, *Cxcl1*, *Lrg1*, *Pfn1*, and *Rnase4*) in these 3 cell types following Tet1 deletion (Figure 5I/5J/5L, Tables S5 and S6). In AT2 cells specifically, 38 DEGs were upregulated while 40 DEGs were downregulated in KO-HDM compared to WT-HDM. Among these 78 genes, several have previously been linked to asthma, including *H3f3b* (Fig 5M) ^43^, *Cd200 (*Fig 5N) ^63^, *Phlda1* (Fig 5O) ^64^, and *Acot7* ^65^ (Figure 5 and Table S6). Further, we identified 32 significantly enriched pathways in the DEGs (p-value<0.05) from AT2 cells (Table S7), including stress response pathways [Unfolded protein response/Endoplasmic Reticulum Stress Pathway/ Protein Ubiquitination Pathway/NRF2-mediated Oxidative Stress Response, Peroxidation of lipid (Figure 6E)], in which most genes showed downregulation in KO-HDM. In particular, Unfolded protein response was predicted to be inhibited (z-score= -2.236). Similar to the analysis in all cells, several genes encoding chemokines and cytokines (e.g., *Ccl20*, *Cxcl6*, *Cxcl5*, *Cxcl3*, and *Cxcl1*) showed reduced expression in KO-HDM (Table S6). Interestingly, *Sftpc* was significantly downregulated in AP and BAS cells in KO-HDM. This gene was downregulated in AT2 cells in KO-HDM compared to WT-HDM (FDR=6.75E-20, fold of change=1.09), and in KO-Saline compared to WT-Saline (FDR=4.01E-49, fold of change=1.05) but did not meet out our fold of change cutoff. Surfactant protein C (SP-C) encoded by this gene enhances the ability of surfactant phospholipids to decrease alveolar surface tension ^66^ and the deficiency of SP-C promotes alveolar instability, associated with lung inflammation and injury ^67 68^. In summary, our data suggest that following HDM challenges, loss of Tet1 led to downregulation of genes in stress response pathways and surfactant proteins, which may lead to significant cell apoptosis and tissue injury (suggested by upregulation of genes involved in tissue damage and repair) and additional lung inflammation and remodeling.

### Tet1 knockout and HDM exposure induced cell type-specific changes in transcription factor (TF) activity in the EpCAM^+^ lung epithelial cells

As Tet1 may regulate gene expression through interacting with TFs, TF activity in individual cell types was explored using DoRothEA ^20–22^. Consistent with previous findings ^69–71^, we observed cell type-specific TF activity, consistent with their cell-type specific regulation of gene expression programs (Figure 7A). For example, *E2f1-4* and *Tfdp1* (both promotes expression of cell-cycle related network ^69^) were highly activated in BAS compared to all other cell types, while the activities of *Rest* (a repressor of neuronal genes in non-neuronal cells ^72^) and *Rreb1* were lowest in Neuroendocrine cells. Additionally, there was relatively high activity of *Rfx1* (promotes ciliogenesis ^73, 74^) in ciliated cells. Meanwhile, because AT2 cells were the most abundant cell type, there were generally only small changes in TFs activity compared to the average level across all cells in our heatmap (Figure 7A). However, a group of TFs, including *Foxa1*, *Sox10*, and *Runx2*, showed relatively high activity levels in AT2 cells, and these TFs are essential in conducting airway and alveolar epithelium development and repair ^75–77^. Subsequently, we found that TF activity levels were altered by HDM and Tet1 loss among individual cell types, such as AT2 cells (Figure 7B-7E) and ciliated cells (Figure 7F-7H). For example, regardless of Tet1 status, *Foxa1* (also decreased in expression in HDM samples in AT2 cells, Table S5) and *Hnf1a* activities were reduced by HDM in AT2 cells (Figure 7B), while the activities of *Rest, Rreb1*, and *Nr2c2* were increased by HDM in both AT2 cells and ciliated cells (Figure 7B and 7F). Meanwhile, *Smad3*, *Smad4*, and *Sp3* activities were decreased by HDM in ciliated cells (Figure 7F). When specifically looking for effects of Tet1, we found that the activities of *Myc, Rest* and *Rreb1* were consistently reduced when Tet1 was lost in AT2 cells regardless of HDM status (Figure 7C). Hif1α, whose expression is regulated by Tet1 ^78^ and may also regulate Tet1 expression ^79^, showed reduced activity in KO-Saline (Figure 7D). Increased *Foxa1* activity was only observed in KO-HDM compared to WT-HDM (Figure 7E). In ciliated cells, however, nearly all effects of Tet1 loss were specific to treatment state (Figure 7F). With saline treatment, activities of *Nfe2l2 (*encoding Nrf2), *Nfe2l1* (encoding Nrf1) and their binding partner Mafk (Figure 7G) ^80^ were reduced in TET1 KO cells compared to wildtype cells, and these TFs are involved in various stress responses including oxidative stress ^81, 82^ and regulate the expression of ROS-detoxifying enzymes including *Gstp1*, *Gstm1* and *Gpx2* ^83^. Meanwhile, TFs involved in the aryl hydrocarbon receptor signaling pathway (AhR) and the TGFβ signaling pathway (Smad3, Smad4) were higher in KO-HDM compared to WT-HDM, even though HDM challenges reduced their activity (Figure 7F/7H). Collectively, our data suggest that TFs respond to HDM challenges and Tet1 loss in a cell type specific manner, which may contribute to the cell-type specific transcriptomic changes and promote lung inflammation.

**Figure 7.**
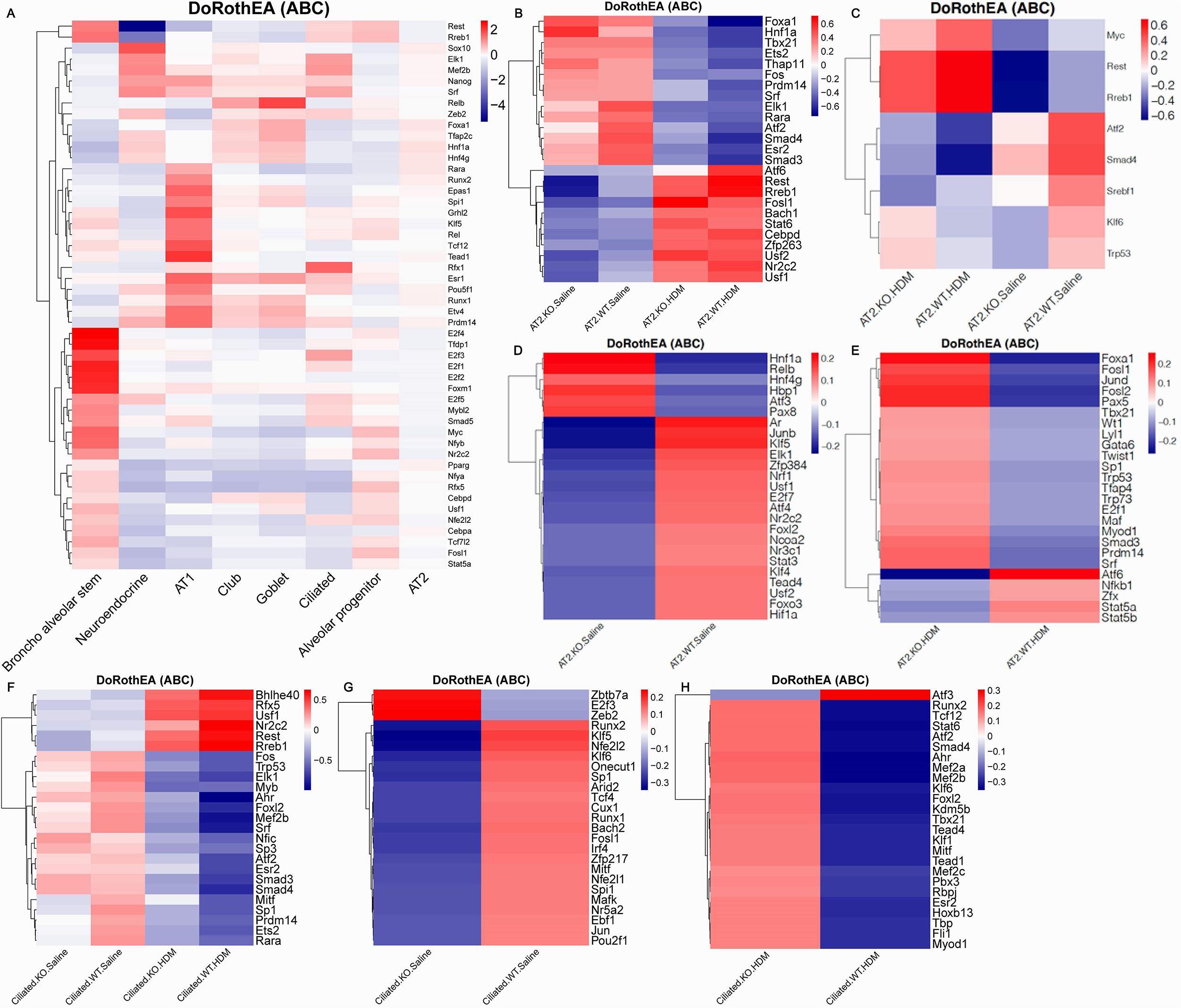
Heatmaps of transcription factor (TFs) activities analyzed using DoRothEA. (A) Top 50 most variable TFs by gene activity across eight individual cell types. (B) Top 25 TFs across all AT2 cells. (C) TFs that were in the top 25 in both the KO-HDM vs WT-HDM and KO-Saline vs WT-Saline comparisons in AT2 cells. (D) Top 25 TFs in the KO-Saline vs WT-Saline comparison that were not in the top 25 for KO-HDM vs WT-HDM in AT2 cells. (E) Top 25 TFs in the KO-HDM vs WT-HDM comparison that were not in the top 25 for KO-Saline vs WT-Saline in AT2 cells. (F) Top 25 most variable TFs in ciliated cells. (G) Top 25 most variable TFs in ciliated cells in KO-Saline vs. WT-Saline. (H) Top 25 most variable TFs in ciliated cells in KO-HDM vs. WT-HDM.

### HDM, not Tet1, promotes the differentiation of AP cells and BAS into AT2 cells

As observed above, following HDM challenge, AT2 cells were reduced in prevalence while AP and BAS cells were increased in WT-Sal and KO-Sal mice (Figure 2 and Table S2). It is well established that AT2 cells can differentiate from AP and BAS cells ^84, 85^. Therefore, single-cell trajectory analysis was performed to explore the roles of HDM and/or Tet1 KO in AT2 cell differentiation (Figure S4/S5). Initially, AT2, AP, and BAS cells from all groups were combined (Figure S4A) and then cell fates were separated at branch point 4 (Figure S4B). Our data support that AT2 cells differentiated from AP and BAS cells. Comparing WT-Sal to WT-HDM and KO-Sal to KO-HDM, HDM challenged cells have a different cell type distribution than saline treated cells (Figure S4C-4H). However, this was not observed in the WT-Sal vs KO-Sal comparison, as no clear branch point was identified to separate cell fates (Figure S4I-4K). Thus, the heatmap of this comparison could not be generated (Figure S5). Deletion of Tet1 had no significant effects on the AP AT2 or BAS AT2 trajectories (Figure S4I-4N). Combined with previous evidence in Figure S4 and Table S2, these results suggest that HDM-induced AT2 cells injury promoted the proliferation and differentiation of AP cells and BAS. Meanwhile, Tet1 knockout did not appear to affect the differentiation of AP cells and BAS into AT2 cells.

## DISCUSSION

In this study, we used scRNA-seq to understand the role of Tet1 in EpCAM^+^ CD45^-^ CD31^-^ lung epithelial cells in HDM-induced lung inflammation at the single-cell level. Eight types of lung epithelial cells and one population of stromal cells were identified. Among them, AT2 cells were the most abundant type. HDM challenge increased the percentage of AP, BAS, and goblet cells, and decreased the percentage of AT2 and ciliated cells. Furthermore, we observed cell type-specific transcriptomic alterations in various types of lung epithelial cells following HDM challenge and/or Tet1 deletion in mice. Via bulk cell and cell type-specific pairwise comparisons, we found upregulation of pro-inflammatory cytokines/alarmins and genes involved in tissue damage/repair, and downregulation of genes in multiple oxidative stress response pathways when Tet1 was knocked out with or without HDM challenges. These transcriptomic regulations were accompanied by changes in TF activities. Trajectory analysis also supported that HDM may enhance the differentiation of AP and BAS cells into AT2 cells, independent of Tet1. Taken together, our data suggest that different lung epithelial cells played both common and unique roles in allergic lung inflammation. Tet1 deletion altered the networks of genes regulating oxidative stress responses, alarmins, and tissue injury/remodeling in various lung epithelial cells, with an overall effect of promoting allergy-induced lung inflammation and remodeling in mice (proposed model in Figure 8).

**Figure 8.**
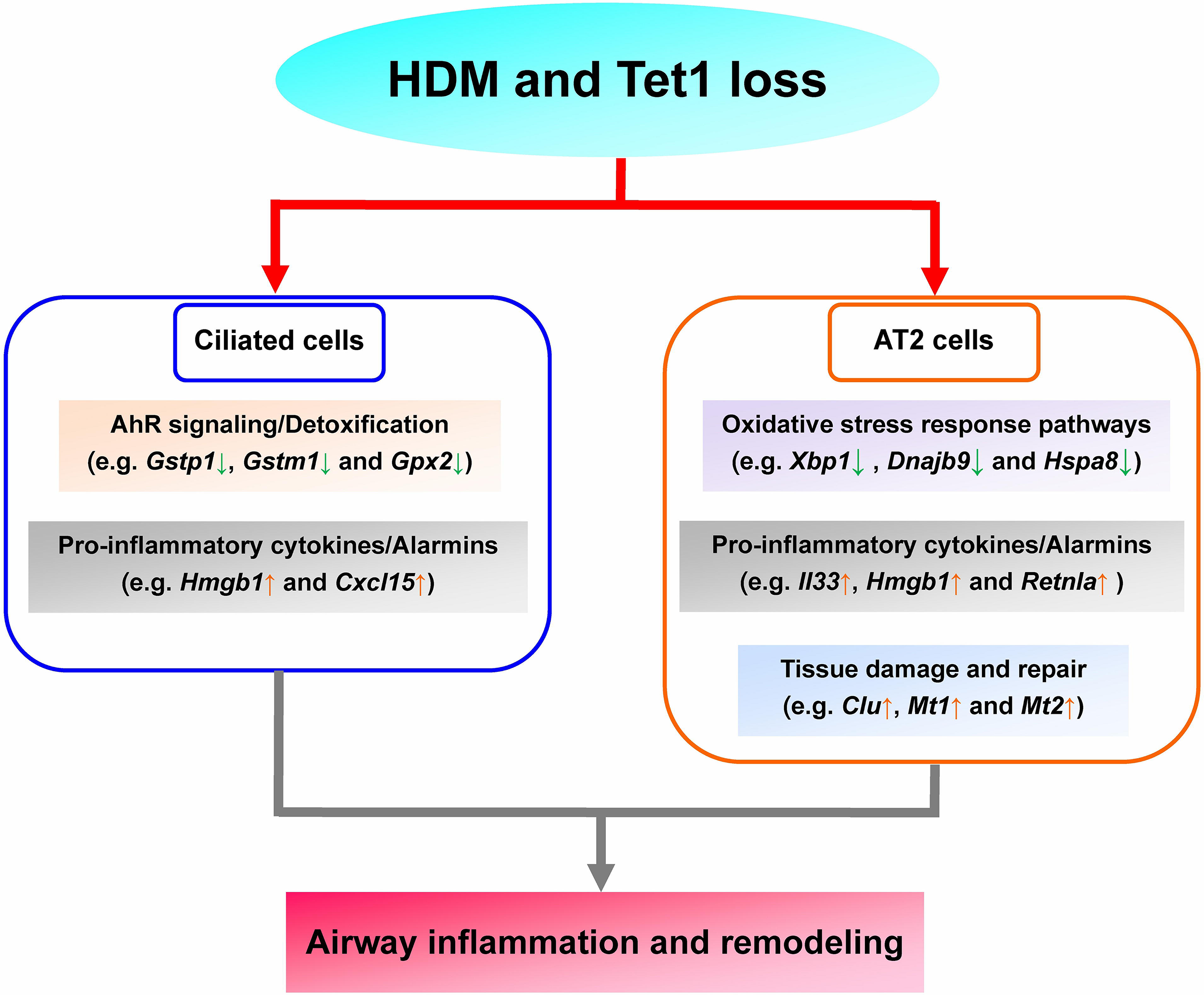
Proposed model. Tet1 deletion promotes allergy-induced lung inflammation and remodeling in mice, involving down-regulation of genes in AhR signaling/detoxification (ciliated cells) and oxidative stress responses (AT2 cells), and up-regulation of genes in pro-inflammatory cytokines/alarmins (AT2 and ciliated cells) and tissue damage and repair (AT2 cells).

The lung is an organ with continuous exposure to environmental challenges, such as allergens and pathogens. The epithelium is the first line of defense for the lung. Lung epithelium (airway epithelium ^86^ and alveolar epithelium ^87, 88^) injury and inflammation play critical roles in asthma. Our previous study ^9^ found that Tet1 deletion enhanced HDM-induced lung inflammation in mice, and that was linked to transcriptomic changes in proinflammatory molecules (*Il33* and *Isg15*) and detoxification enzymes (*Gsto1* and *Aldh1a1*) in total lung RNA. Furthermore, our previous results indicated that 59 pathways, including NRF2-mediated Oxidative Stress Response, AhR Signaling, and the Interferon (IFN) signaling pathway, were significantly altered. However, the role of Tet1 loss in specific lung cell types in allergic airway inflammation and the contribution of individual cell types in HDM-induced whole lung transcriptomic alterations remained unknown. In the present study, we used scRNA-seq to explore transcriptomic changes in different types of lung epithelial cells at the single-level, addressing the aforementioned questions. Two strategies were taken to explore the effects of Tet1 knockout in HDM-induced lung inflammation. First, we searched for DEGs with consistent changes in direction among 3 comparisons (WT-HDM vs WT-Sal, KO-Sal vs WT-Sal, and KO-HDM vs WT-Sal). This set of genes was similarly regulated by HDM challenge, Tet1 KO, and HDM challenge plus Tet1 KO. Using this strategy, we identified a set of genes [Cluster 1 (45 unique genes after removing duplicates, Table S9), which include alarms (*Il33*, *Hmgb1*, *S100a6*, etc.) in AT2 cells and detoxification enzymes (*Gstp1*, *Gstm2* and *Gpx2*) in ciliated cells]. Notably, Tet1 knockout resulted in upregulation of alarms and downregulation of detoxification enzymes (Table S10). Second, we searched for additional changes comparing WT-HDM vs KO-HDM. This strategy identified another set of genes [Cluster 2 (Figure 5, Tables S3 and S5)], including 72 genes from EpCAM^+^ cells and 195 genes from AT2, AP and/or BAS cells (103 unique genes after removing duplicates)]. In this cluster, genes in multiple stress response pathways were downregulated and several genes in tissue repair/remodeling (e.g. *Clu*) and inflammatory response (e.g. *Cd200*) were significantly upregulated (Figure 6E/6F and Tables S6/S11). Not all genes identified from the bulk cell analysis were found in the analyses in individual cell types, and vice versa. Yet in general, several genes involved in oxidative stress responses, pro-inflammatory responses, alarmins, AhR signaling/detoxification, and tissue damage and repair were differentially expressed in our dataset (Figure 8), and many are known to contribute to asthma based on evidence from animal models and studies in human subjects. Our study, for the first time, not only showed cell-specific expression patterns at baseline and following HDM challenges in the lung epithelium, but also revealed a key role for Tet1 to regulate these genes transcriptionally in a cell-type specific fashion and promote HDM-induced airway inflammation. Future studies focusing on mechanisms through which Tet1 regulates these genes in the lung epithelium would provide more clarity on this regulatory process.

AT2 cells were the most abundant cell type in our dataset (69.7% in total). Although no significant pathological alterations in alveoli were observed in either asthmatic patients or animal models, several studies ^87–89^ have reported that AT2 cells are a major contributor of many inflammatory mediators involved in asthma, In particular, Ravanetti L, *et al.* supported that AT2 cells are one major source of IL-33 in mice with HDM-induced lung inflammation ^89^. Our present study supports that HDM may enhance AT2 cells differentiation from AP and BAS cells (Figure S4), which was likely due to HDM-induced AT2 cells injury and death (Figure 2 and Table S2). Our cell type-specific transcriptomic analysis showed that AT2 cells were one of the major sources of a cluster of inflammatory mediators and molecules, especially *Il33* (Fig 4F)*, Hmgb1* (Fig 4J), *Ager* (Fig S3-A1), and *Retnla* (Fig S3-A2), in allergen-induced lung inflammation (Table S6). Allergens may induce lung Th2 inflammation through AT2 cell injury which releases alarms such as *Il33* ^25^ and *Hmgb1* ^90^, and Tet1 played a critical role in this process by regulating their expression. Additionally, when comparing KO-HDM to WT-HDM in AT2 cells, we found that genes involved in tissue damage and repair were upregulated, while genes involved in oxidative stress responses were downregulated; most of these genes were also differentially expressed in EpCAM^+^ cells (Table S6). Downregulation of oxidative stress response pathways, especially unfolded protein response (UPR), may result in apoptosis and cell death ^91^, and the roles of UPR in oxidative stress response, cytokine production and asthma pathogenesis are emerging ^92^. Taken together, our data suggest that while AT2 cells from Tet1 knockout mice at base line (no HDM) have a pro-inflammatory transcriptomic signature (elevated alarms and reduced detoxification enzymes), HDM challenges may further induce cell damage and death due to elevated oxidative stress and downregulated stress responses.

Ciliated cells are another major cell type within airway epithelium. Ciliated cells are not only essential for mucociliary clearance through their motile cilia coordinated directional movement, but can also sense and respond to allergens, air pollution particles, and pathogens ^32, 93^. Asthma can cause cilia dysfunction and ultrastructural abnormalities, such as ciliary beat frequency decreasing, dyskinetic and immotile cilia, and disruption of tight junctions, which were closely associated with asthma severity in patients ^94^. Meanwhile, multiple studies ^35, 95^ suggest that ciliated cells are a source of cytokines, such as IL-33 and TSLP, in asthma. In our single-cell analysis, we only observed a trend of increase for *Il33* at baseline when Tet1 was deficient and no further changes following HDM treatment. However, *in vitro* studies using HBECs showed that loss of Tet1 significantly increased *IL33* expression following 24 hours of HDM challenges (Supplementary Figure S6), which support that Tet1 deficiency promotes the predisposition of allergic inflammation in ciliated cells. Further, we found that *Gstp1*, *Gstm1*, and *Gpx2* (detoxification enzymes that have been associated with asthma) were all downregulated with HDM and/or Tet1 loss in ciliated cells (Figure 4L-4N). Our previous studies in HBECs showed the regulation of detoxification enzymes by Tet1 in HEBCs as well ^9^. Together, these data highlight the unique role of ciliated cells and suggest that Tet1 regulates HDM-induced lung inflammation via regulation of AhR signaling/Detoxification and Pro-inflammatory cytokines/Alarmins in ciliated cells (Figures 6D, 7F and 8).

Since we previously observed significant increases in AHR and BALF eosinophils, as well as significant gene expression differences in bulk lung RNA from KO-HDM mice and WT-HDM mice ^9^, transcriptomic differences between KO-HDM and WT-HDM cells were further explored. We identified 72 DEGs, associated with 29 significantly enriched pathways, between KO-HDM and WT-HDM EpCAM+ cells. A total of 195 cell-type specific DEGs were found, including 78 DEGs in AT2 cells. However, no overlapping DEGs with consistent directions of change among all 4 comparisons (WT-HDM vs WT-Sal, KO-Sal vs WT-Sal, KO-HDM vs WT-Sal, and KO-HDM vs WT-HDM) were observed in either all EpCAM^+^ cells together or in individual cell types. Additionally, we also noted that some DEGs identified from bulk RNA in our previous study, such as *Il13*, *Muc5ac*, *Cyp1a1*, and *Irf7*, were not differentially expressed in our present study. We sometimes observed opposite effects of Tet1 knockout following HDM treatment to saline treatment compared to our previous bulk RNA analysis (e.g. *Scd1* and *Pfn1* in AT2 cells). This inconsistency may be because HDM-induced transcriptomic changes masked the effects of Tet1 knockout in lung epithelial cells. Meanwhile, increased noise in scRNA-seq data, lower power, or the differences in how DEGs were defined in the scRNA-seq dataset compared to our previous bulk RNA-seq studies were also potential reasons for these inconsistencies. Another possible explanation is that only lung epithelial cells (EpCAM^+^ cells) were analyzed. These DEGs identified from bulk RNA-seq may have altered expression in other lung cell types such as inflammatory cells, airway smooth muscles, and vascular endothelial cells, which will be pursued in future studies. In the case of Muc5ac, we found that it is predominantly expressed in goblet cells and induced by HDM challenges in both WT (p=0.017, fold of change=6.12) and KO mice (p=0.00014, fold of change=3.62). However, due to the small number of goblet cells, none of these comparisons reached FDR significance and no additional significant changes were observed comparing WT-HDM and KO-HDM at p value<0.05. Appreciable transcriptomic changes were also observed in other types of lung epithelial cells, including AP cells, BAS cells, club cells, goblet cells, and neuroendocrine cells, following HDM exposure and Tet1 deletion in mice. However, no transcriptomic differences in AT1 cells were found, likely due to the limited number of AT1 cells we isolated from the lungs (Table 1). Interestingly, we also observed downregulation of several asthma-associated genes (e.g. *Atf3* ^96^ and *Btg2* ^97^) that are normally increased by HDM challenges, which is possibly due to the time-dependent expression pattern of these molecules during acute phase of inflammation and needs further investigation. Nevertheless, our analysis revealed novel changes that were not found in our previous bulk RNA-seq analysis, which shed light on possible mechanisms of Tet1 in regulating airway inflammation in the presence of HDM.

Additionally, our data also suggest that HDM and/or Tet1 KO in mice lungs led to cell type-specific changes in TF activity levels (Figure 7). For example, *Foxa1* had reduced activity following HDM treatment in AT2 cells, but activity was higher in the KO-HDM samples compared to WT-HDM samples (Figure 7B). Foxa1 has been shown to play a role in maintaining the integrity of epithelial barriers in primary human bronchial epithelial cells ^77^, so changes in Foxa1 activity could impact the airway epithelium. *Foxa1* also had significantly decreased expression in HDM samples compared to saline samples in AT2 cells (Table S5). Additionally, AhR (aryl hydrocarbon receptor) showed decreased activity levels in HDM samples compared to saline samples in ciliated cells (similar between KO-saline and WT-saline), but KO-HDM samples had higher AhR activity than WT-HDM samples (Figure 7F/7H). In our prior study using human bronchial epithelial cells, we hypothesized that Tet1 protects against HDM-induced allergic inflammation at least partially by regulating the AhR pathway ^9^. Data from the current study in ciliated cells supports that AhR activity and the AhR pathway were altered following Tet1 KO and/or HDM treatment (Figure 7F/7H and Figure 8). Meanwhile, the Nrf1/2 activity was lower in KO-Saline mice compared to WT-Saline, which may explain the lower expression of *Gstp1*, *Gstm1* and *Gpx2* in KO-Saline compared to WT-Saline as the Nrf2-mediated pathway is an alternative to the AhR pathway that regulates the expression of antioxidant enzymes ^83^. Smad3, Smad4, Fos, and Atf2 activity levels were all altered by HDM treatment and/or Tet1 KO in AT2 and ciliated cells (Figure 7B/7F/7H). *Fos* also had significantly decreased expression following HDM treatment in AT2 and ciliated cells (Table S5), but not following Tet1 KO. These transcription factors are all part of the AP-1 signaling pathway, which has been previously associated with regulation of asthmatic inflammation ^98, 99^. These activity levels for the transcription factors were estimated using gene expression profiles of their targets, not based on the direct expression of the transcription factor itself. Even if the direct expression of a transcription factor was not significantly altered, it is still possible that the way that the transcription factors are regulating gene expression could be altered by Tet1 KO or HDM treatment, especially considering that Tet1 is known to directly interact with transcription factors ^100, 101^. Taken together, these results indicate that Tet1 plays an important role in regulating TF activity that directly impacts allergen-induced lung inflammation.

This study had several strengths. First, single-cell RNA analysis was used to explore the transcriptome profile of purified lung epithelial cells in allergic inflammation at the single-cell level, which elucidated the specific roles of each epithelial cell type. Second, Tet1^-/-^ mice were used, which is more accurate and reliable than other knockdown methods, such as plasmid transfection, adeno-associated virus (AAV) transfection, etc. Meanwhile, *in vitro* studies were performed to validate our findings in mice. One weakness of this study was a small sample size (two mice in each group, with decent number of cells sequenced within each group). The contributions of other lung cells, such as airway smooth muscle, vascular endothelial cells and immune cells remain unclear in these pathological processes. Future studies will be designed to investigate the therapeutic potential of targeting Tet1 in asthma.

## CONCLUSIONS

Collectively, our results revealed possible mechanisms underlying the protective role of Tet1 in HDM-induced lung inflammation at the single-cell level. Different epithelial cells played common and unique roles in allergic lung inflammation. The major cellular contributors of multiple asthma-associated inflammatory mediators, such as *Il33*, *Hmgb1*, *Retnla*, and *Ager* were identified, and a cluster of novel allergy-associated molecules, such as *Chia1*, *Nnat*, and *Nr4a1* were found in different lung epithelial cells. Future studies should attempt to elucidate the role of these novel targets in asthma. Our results suggest that AT2 cells are essential for Th2 inflammation through stress responses, alarms, tissue injury and repair in allergen-induced lung inflammation, and that these genes are regulated by Tet1. In ciliated cells, Tet1 promotes the Xenobiotic Metabolism AhR signaling pathway, especially the detoxification enzymes, to protect from allergen-induced lung inflammation. Overall, these novel findings support that Tet1 is a potential therapeutic target of asthma.

## Authors’ Contributions

HJ conceived the study. TZ drafted the manuscript with the help of APB and HJ. TZ established and characterized the mouse model with the help of LC. APB performed scRNA-seq data processing and analysis. LC performed sample processing and assisted in library preparation. GQ assisted in scRNA-seq data processing and provided input in cell clustering. All authors have approved the final version of this manuscript. Part of this study has been published as an abstract (https://doi.org/10.1164/ajrccm-conference.2019.199.1_MeetingAbstracts.A6041 and https://doi.org/10.1164/ajrccm-conference.2020.201.1_MeetingAbstracts.A7402).

## Acknowledgements

This work was supported by NIH/NIAID R01AI141569-1A1 (HJ). HJ was also supported by NIH/NIEHS P30ES023513-supported EHSC scholar fund and UC Davis Faculty Startup fund. Funding was also provided by NSF CAREER award 1846559 to GQ.

