## Supplementary figures and images for "Single-cell RNA-seq analysis reveals lung epithelial cell-specific contributions of Tet1 to allergic inflammation"

### Supplementary Figure 1

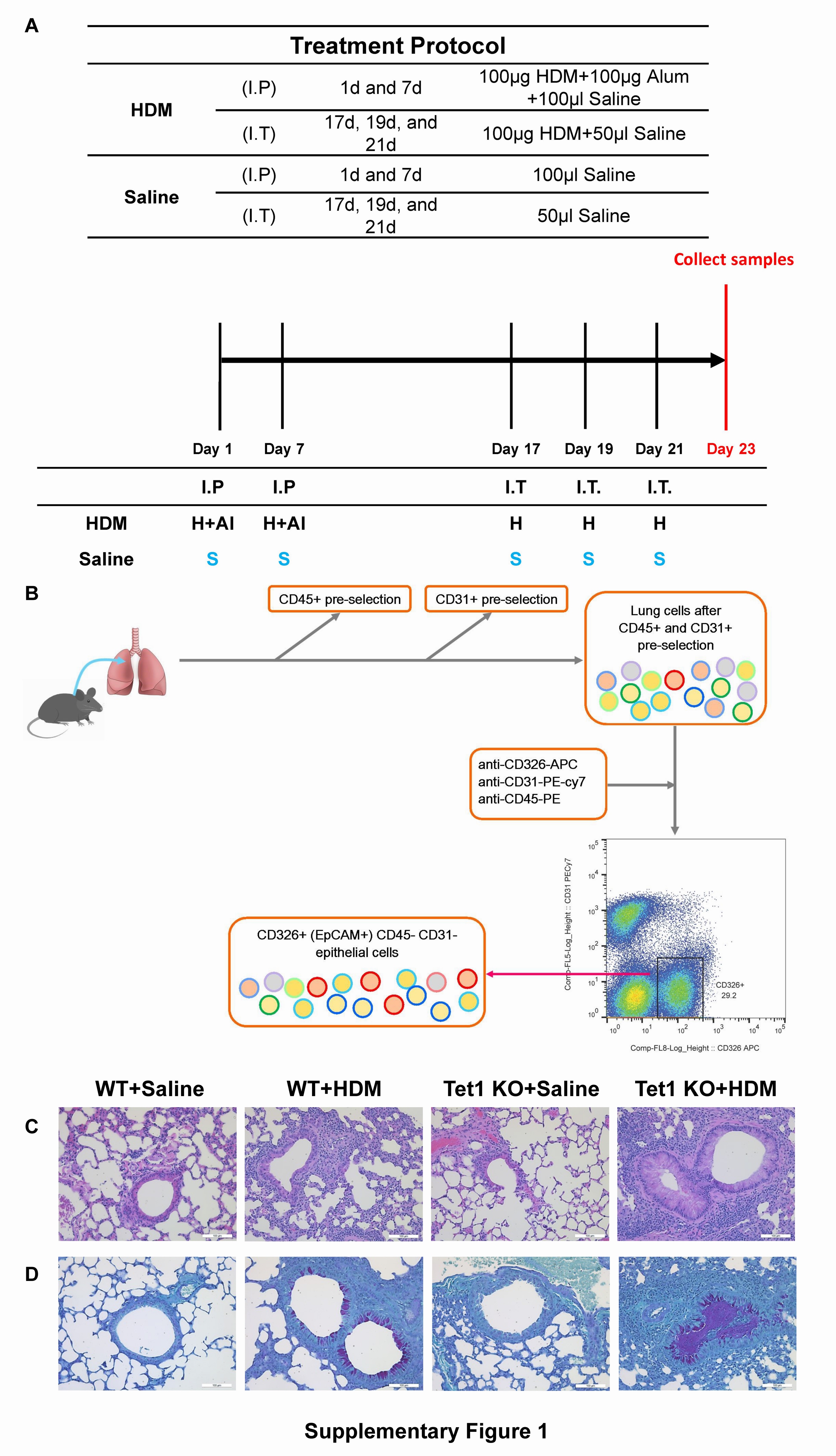

### Supplementary Figure 2

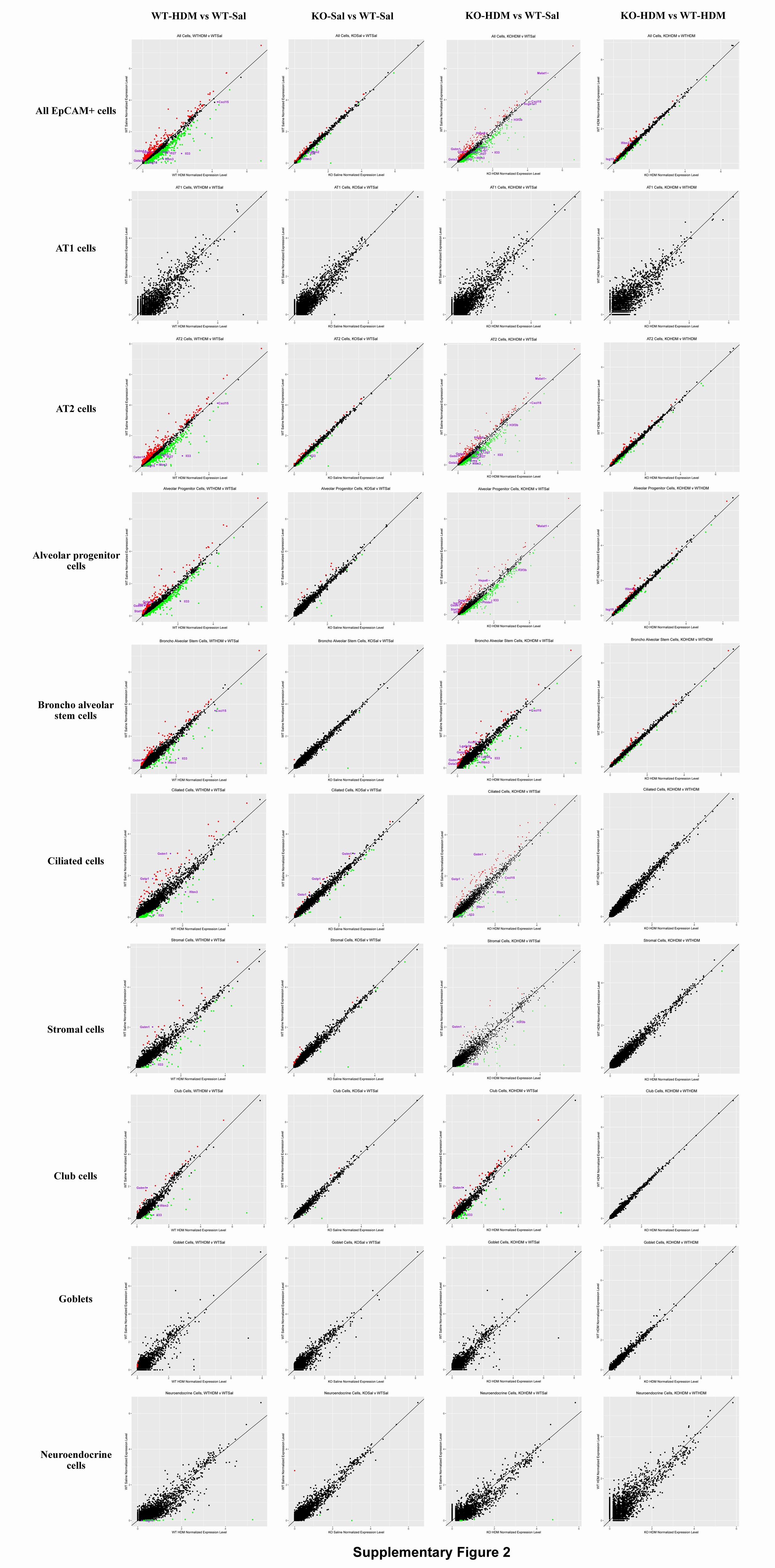

### Supplementary Figure 3

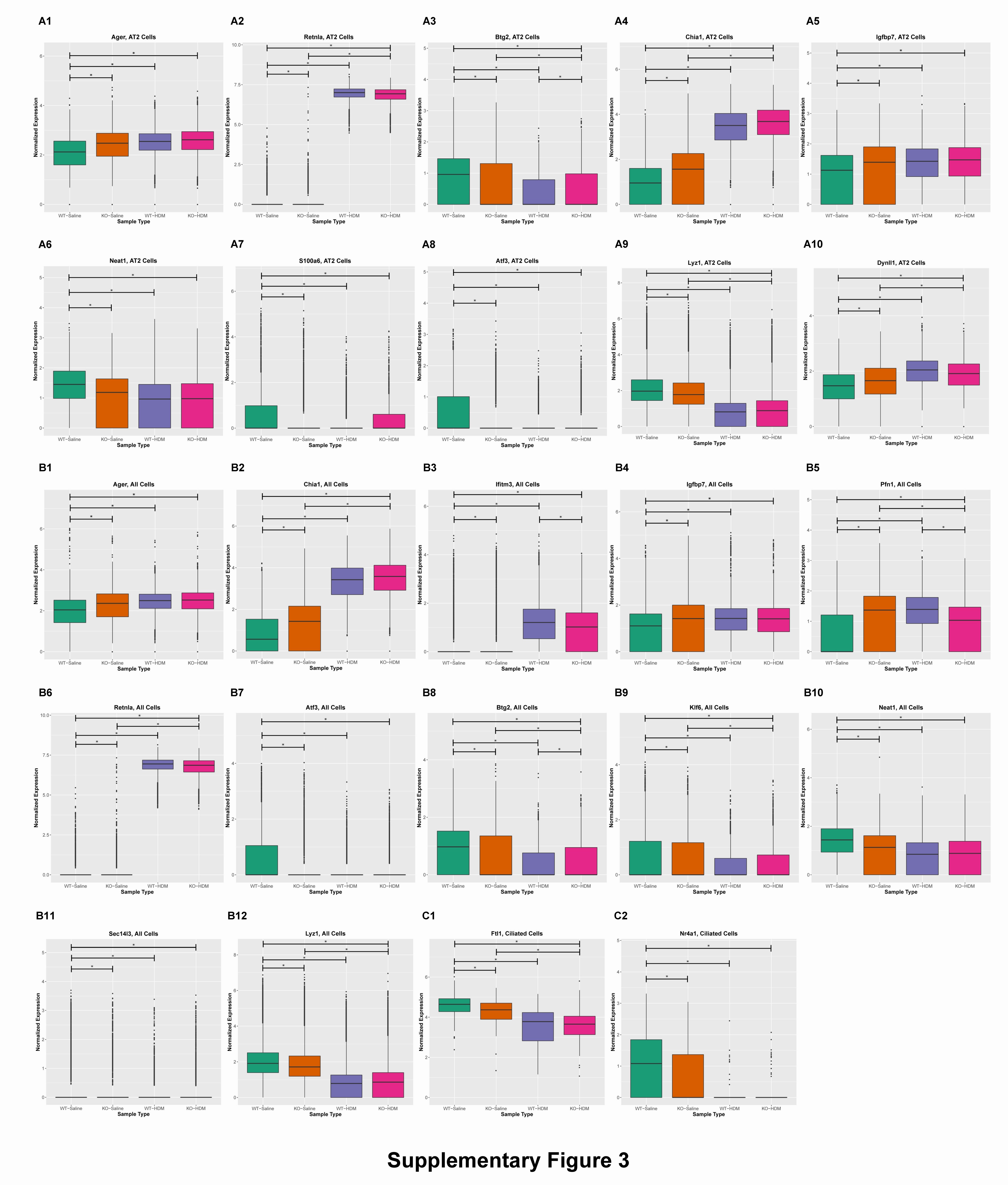

### Supplementary Figure 4

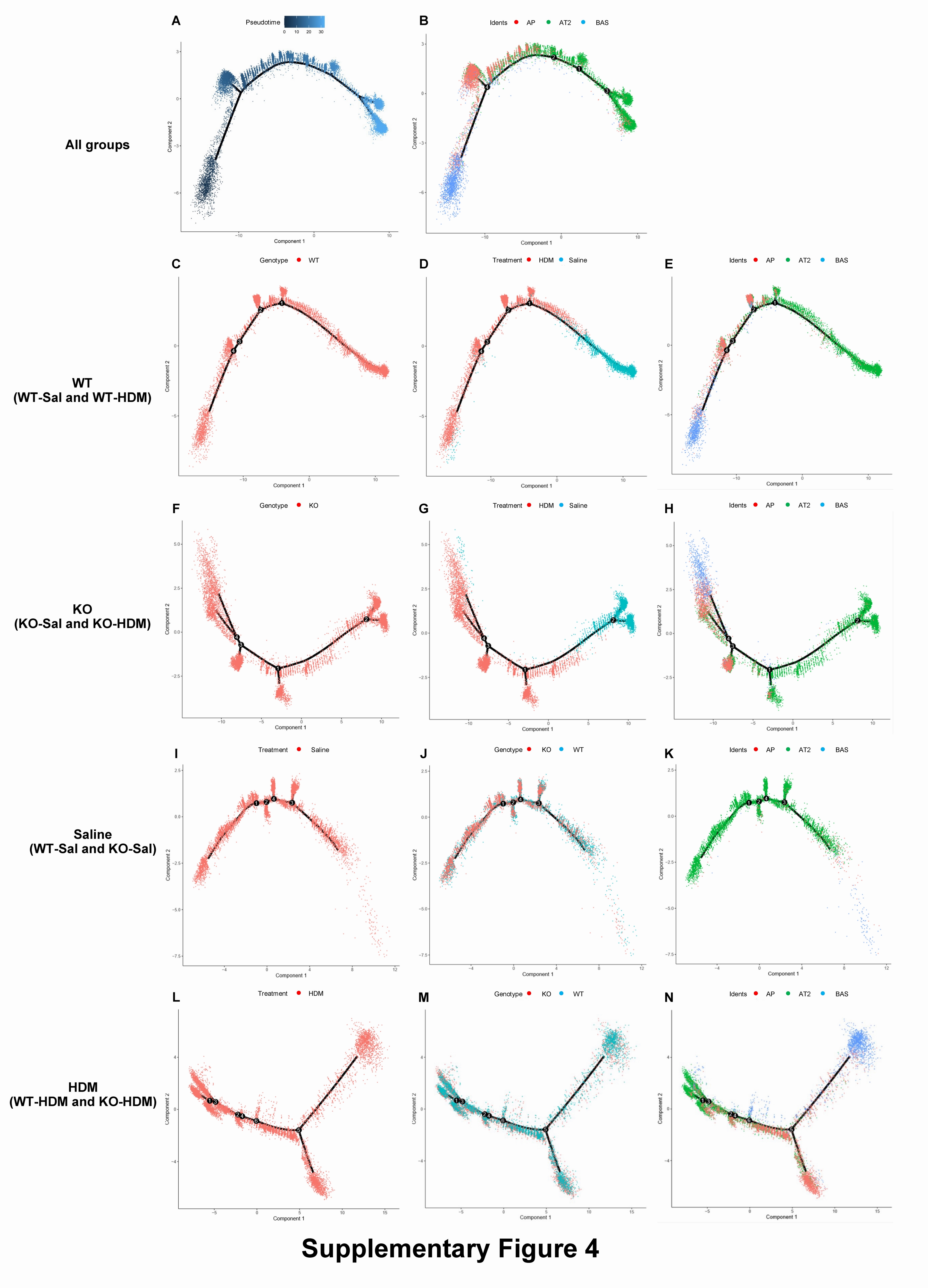

### Supplementary Figure 5

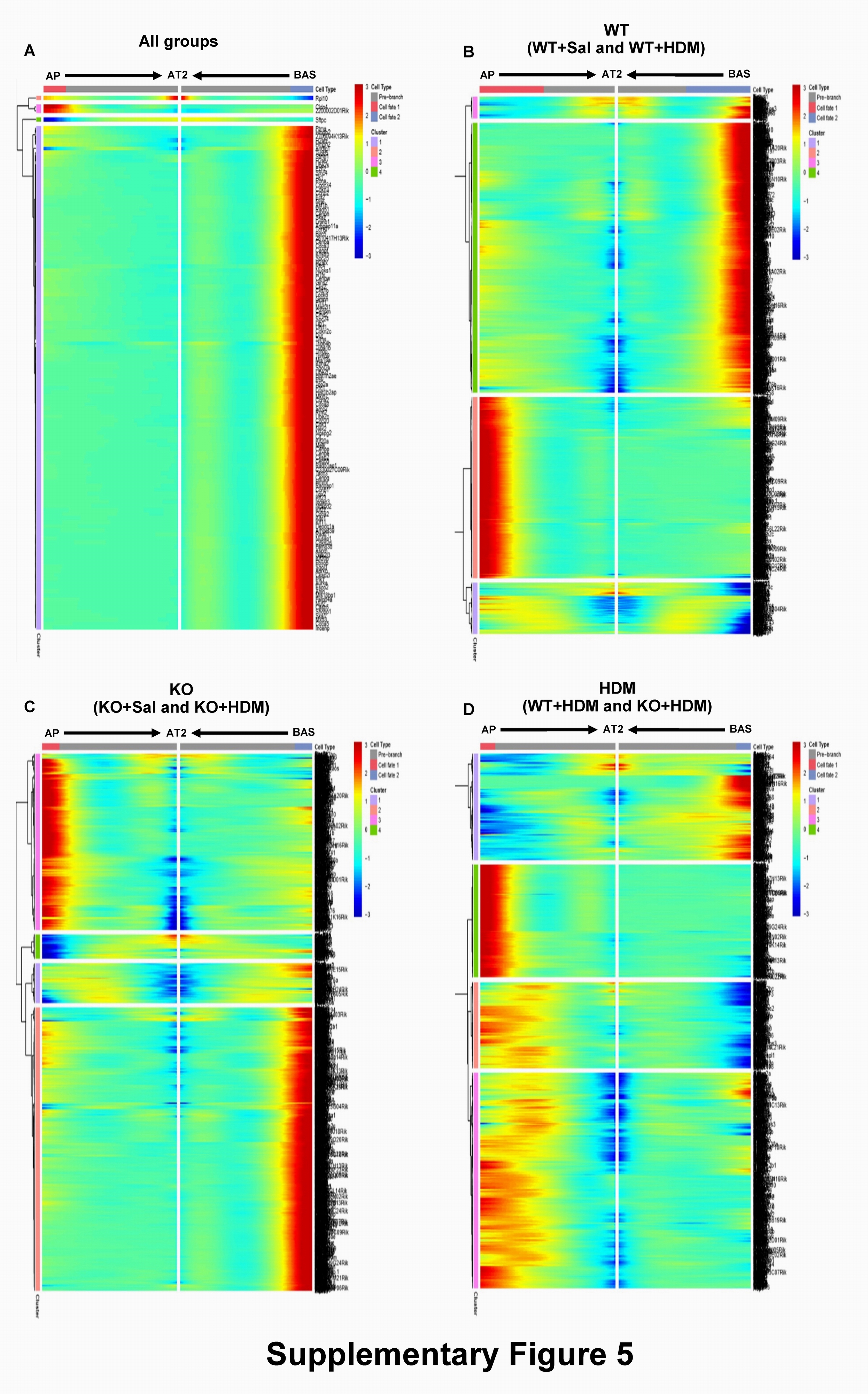

### Supplementary Figure 6

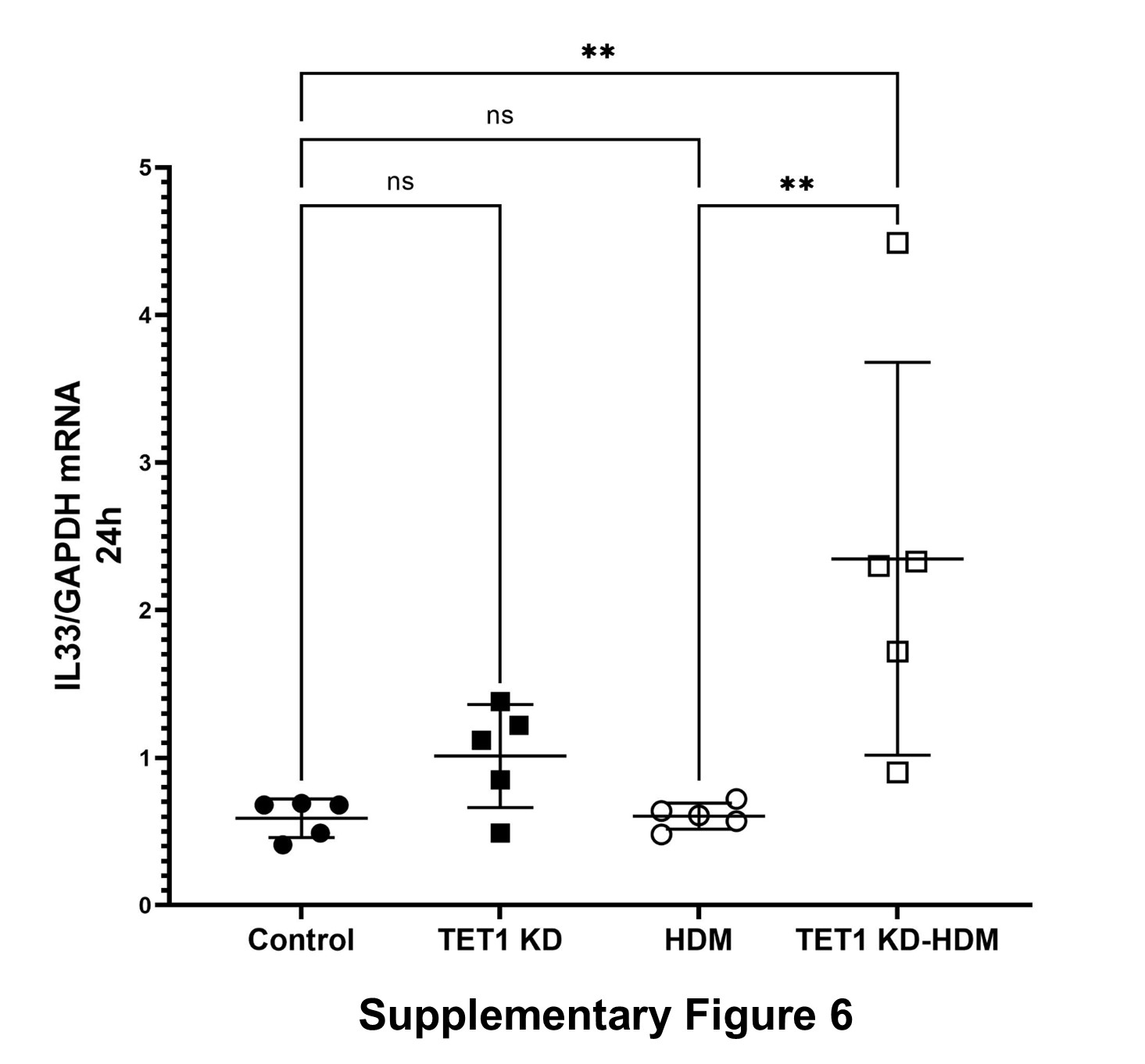
