## Supplementary Information for "Single-cell RNA-seq analysis reveals lung epithelial cell-specific contributions of Tet1 to allergic inflammation"

**Supplementary Methods**

**Animal**

In this study, both Tet1$^{-/-}$ mice (knockout/KO mice) and their littermate Tet1$^{+/+}$ mice (wildtype/WT mice) were used. The establishment of Tet1$^{-/-}$ mouse was described in a previous study [9]. Meanwhile, all procedures on animals were approved by the Animal Experimental Ethics Committee of Cincinnati Children's Hospital Medical Center and University of California, Davis. This study was performed based on the recommendations in the Guide for the Care and Use of Laboratory Animals. All surgery was performed under anesthesia, and all efforts were made to minimize suffering. Specific pathogen free (SPF) mice (16-24g, age 8-12 weeks), were maintained under SPF conditions in the animal facilities. The mice were fed in a temperature-controlled room (12-h dark and light cycles) and offered *ad libitum* access to food and water. Animals underwent an acclimatization period of at least 1 week before the study.

**Murine model of HDM-induced allergic airway inflammation**

In this study, the murine asthma model was induced by HDM as we previously described [9] (Figure S1A). Briefly, the mice were divided into 4 groups, with 2 mice in each group: WT-Saline, WT-HDM, KO-Saline, and KO-HDM. Mice in the WT-HDM and KO-HDM groups were challenged with two intraperitoneal (I.P) injections of 100μg HDM [including 61.3μg of protein, Dermatophagoides pteronyssinus, Greer Laboratories (Lenoir, NC)] plus 100μg aluminium (Alum) in 100μl Saline followed by three intratracheal (I.T) instillations of 100μg HDM in 50μl Saline. Meanwhile, mice in the WT-Saline and KO-Saline groups were treated with 50μl Saline instead of HDM. To ensure the successful establishment of the mouse model, the left caudal lung of each mouse was fixed in 10% formalin, embedded in paraffin, and cut into 5μm sections. Hematoxylin and eosin (H&E) staining was performed to observe pulmonary pathological alterations. Periodic acid–Schiff (PAS) staining was used to observe goblet cell hyperplasia and airway mucus secretion.

'.

**Ingenuity pathway analysis** In the current study, canonical pathway analysis and diseases and functions were carried out using the Ingenuity Pathway Analysis (IPA, Ingenuity Systems, Redwood City, CA). Meanwhile, statistical significance was set at $p<0.05$ (−log (P-value) >1.3).

**HBECs growth and TET1 knockdown**

HBEC were grown in 1X K-SFM supplemented with EGF, pituitary extract, and pen-strep (complete media). A 12 well plate was seeded with cultured HBEC with near confluent numbers and incubated overnight for attachment. After 24 hours, the complete media was replaced with 1X K-SFM with no serum supplements (minimal media) and incubated for an additional 8 hours. TET1 was knocked down by adding 30 pmol/mL siRNA (ThermoFisher) and incubated for 24 hours in minimal media. House dust mite (HDM, Greer) was added to HBEC at a concentration of 25ug/mL and incubated for another 24 hours. Cells were then collected for library preparation and RNA extraction for analysis. There were four different sample types in our study: 1) wildtype (WT) saline, 2) WT house dust mite (HDM), 3) TET1 knockdown (KD) saline, and 4) KD HDM.

**RT-qPCR analysis**

Total RNA was reverse-transcribed to cDNA using the VILO cDNA kit (Thermo Fisher Scientific, Waltham, MA, USA) according to the manufacturer's instructions. Real-time quantitative PCR was performed using the SYBR Green Master Kit and Quantstudio instrument. PCR was carried out in duplicates for each fraction and the mean Ct value of the triplicate reaction was normalized against the mean Ct value of GAPDH. IL33 primers: Forward, 5'-CCT GTC AAC AGC AGT CTA C-3'; Reverse, 5'-TTG GCA TGC AAC CAG AAG TC-3'.

**Supplementary Figure Legends**

**Supplementary Figure S1.**

(A) Treatment Protocol;

(B) Lung epithelial cells ($CD326^+$ $CD31^-$ $CD45^-$ cells) sorting. Representative sorting images (one from WT and one from KO are shown);

(C) and (D) Lung pathological alterations and airway mucus secretion in HDM-induced airway inflammation in mice. [(C) HE staining. The figure demonstrates a representative view (×200) from each group. (D) PAS staining. The figure demonstrates a representative view (×200) from each group.]

**Supplementary Figure S2.** qq plots (WT-Sal vs WT-HDM, WT-Sal vs KO-Sal, KO-HDM vs WT-sal, and KO-HDM vs WT-HDM) for DEGs in all $EpCAM^+$ lung epithelial cells and 9 individual cell types. Differentially expressed genes (FDR<0.05) are either marked in red or green. Individual genes mentioned in the text are highlighted in purple.

**Supplementary Figure S3.** The expressions of asthma-associated DEGs (WT-Sal vs WT-HDM, WT-Sal vs KO-Sal, and KO-HDM vs WT-Sal overlapped DEGs) in (A1-12) $EpCAM^+$ lung epithelial cells, (B1-10) AT2 cells, and (C1-2) ciliated cells.

**Supplementary Figure S4.** Single-cell trajectory analysis reveals the differentiation of AP and BAS cells into AT2 cells.

(A) and (B) Single-cell trajectory of AP, BAS, and AT2 cells in all mice.

(C), (D), and (E) Single-cell trajectory of AP, BAS, and AT2 cells in WT mice with saline treatment and HDM challenge.

(F), (G), and (H) Single-cell trajectory of AP, BAS, and AT2 cells in Tet1 KO mice with saline treatment and HDM challenge.

(I), (J), and (K) Single-cell trajectory of AP, BAS, and AT2 cells in WT and Tet1 KO mice with saline treatment.

(L), (M), and (N) Single-cell trajectory of AP, BAS, and AT2 cells in WT and Tet1 KO mice with HDM challenge.

**Supplementary Figure S5.** Expression heatmap of dynamically expressed genes ordered across single-cell trajectory analysis reveals the differentiation of AP cells and BAS into AT2 cells.

(A) Heatmap depicting genes with a branch-dependent manner for branch point 4 in all mice.

(B) Heatmap depicting genes with a branch-dependent manner for branch point 4 in WT mice with saline and HDM challenge.

(C) Heatmap depicting genes with a branch-dependent manner for branch point 3 in Tet1 KO mice with saline and HDM challenge.

(D) Heatmap depicting genes with a branch-dependent manner for branch point 5 in WT mice and Tet1 KO mice with HDM challenge.

Color scheme represents the z-score distribution from -3.0 (blue) to +3.0 (red). Each row represents the dynamic expression of a gene. Genes along the differentiation of AP cells and BAS into AT2 cells are clustered into 4 blocks. Cell types: Grey: AT2 cells, Red: AP cells, Blue: BAS.

**Supplementary Figure S6.** Impact of TET1 knockdown and HDM treatment in Il33 expression in HBECs.

* represents P<0.05; ** represents P<0.01; ns represents non significant.

**Supplementary Tables**

**Supplementary Table S1.** Markers used to define all the cell types.

**Supplementary Table S2.** Cell number in each group identified by scRNA-seq analysis.

**Supplementary Table S3.** Differentially expressed genes (DEGs) from bulk analysis (all $EpCAM^+$ cells).

**Supplementary Table S4.** Enriched pathways with DEGs from bulk analysis by IPA analysis.

**Supplementary Table S5.** Differentially expressed genes from each cell type.

**Supplementary Table S6.** Examples of cell type-specific changes induced by HDM challenges and Tet1 loss.

"+" and "-" mean genes that were increased and decreased with a false discovery rate (FDR) ≤0.05 and a fold change ≥1.2.

N.S. (non-significant) means differential genes without a false discovery rate (FDR) ≤0.05 and/or a fold change ≥1.2.

N.E. (no-expression) means genes weren't detectable.

**Supplementary Table S7.** Enriched pathways with AT2-specific DEGs by IPA analysis.

**Supplementary Table S8.** Enriched pathways with ciliated cell-specific DEGs by IPA analysis.

**Supplementary Table S9.** Genes with the same directions of changes among WT-HDM vsWT-Sal, KO-SalvsWT-Sal, and KO-HDMvsWT-Sal.

**Supplementary Table S10** Enriched pathways within genes with the same directions of changes among WT-HDMvsWT-Sal, KO-SalvsWT-Sal, and KO-HDMvsWT-Sal in all $EpCAM^+$ cells, AT2 cells, AP, BAS, and ciliated cells.

**Supplementary Table S11**. Enriched pathways within combined DEGs from all $EpCAM^+$ cells, AT2 cells, AP, and BAS with the same directions of changes in KO-HDMvsWT-HDM.
