## Supplementary Table 2 for "Single-cell RNA-seq analysis reveals lung epithelial cell-specific contributions of Tet1 to allergic inflammation"

**Supplementary Table 2. Cell number in each group identified by scRNA-seq analysis**

|  | WT-Sal | WT-HDM | KO-Sal | KO-HDM | Total |
| --- | --- | --- | --- | --- | --- |
| EpCAM^+^ cells | 6378 | 6925 | 5420 | 6714 | 25437 |
| AT1 cells | 18  (0.282%) | 5  (0.072%) | 9  (0.166%) | 8  (0.119%) | 40  (0.157%) |
| AT2 cells | 5502  (86.265%) | 3885  (56.101%) | 4572  (84.354%) | 3770  (56.151%) | 17729  (69.698%) |
| Alveolar progenitor cells | 157  (2.462%) | 1758  (25.386%) | 84  (1.550%) | 1905  (28.374%) | 3904  (15.348%) |
| Broncho alveolar stem cells | 55  (0.862%) | 985  (14.224%) | 62  (1.144%) | 671  (9.994%) | 1773  (6.970%) |
| Ciliated cells | 362  (5.676%) | 66  (0.953%) | 189  (3.487%) | 131  (1.951%) | 748  (2.941%) |
| Stromal cells | 105  (1.646%) | 88  (1.271%) | 354  (6.531%) | 38  (0.566%) | 585  (2.300%) |
| Club cells | 95  (1.489%) | 75  (1.083%) | 115  (2.122%) | 139  (2.070%) | 424  (1.667%) |
| Goblet cells | 5  (0.078%) | 56  (0.809%) | 20  (0.369%) | 45  (0.670%) | 126  (0.495%) |
| Neuroendocrine cells | 79  (1.239%) | 7  (0.101%) | 15  (0.277%) | 7  (0.104%) | 108  (0.425%) |
